# Spatial profiling of microbial communities by sequential FISH with error-robust encoding

**DOI:** 10.1101/2021.05.27.445923

**Authors:** Zhaohui Cao, Wenlong Zuo, Lanxiang Wang, Junyu Chen, Zepeng Qu, Fan Jin, Lei Dai

## Abstract

Spatial analysis of microbiomes at single cell resolution with high multiplexity and accuracy has remained challenging. Here we present spatial profiling of a microbiome using sequential error-robust fluorescence *in situ* hybridization (SEER-FISH), a highly multiplexed and accurate imaging method that allows mapping of microbial communities at micron-scale. We show that multiplexity of RNA profiling in microbiomes can be increased significantly by sequential rounds of probe hybridization and dissociation. Combined with error-correction strategies, we demonstrate that SEER-FISH enables accurate taxonomic identification in complex microbial communities. Using microbial communities composed of diverse bacterial taxa isolated from plant rhizospheres, we show that SEER-FISH can quantify the abundance of each taxon and map microbial biogeography on roots. SEER-FISH should enable accurate spatial profiling of the ecology and function of complex microbial communities.

## Introduction

Spatial structure of microbial communities has been observed across different habitats, ranging from marine biofilms^1^, human gastrointestinal tracts and oral cavities^2, 3^, to plant rhizospheres^4^. For example, microbial localization and density vary widely in animal guts (along both longitudinal and transverse axes), as well as in plant compartments, due to spatial heterogeneity in chemical and oxygen gradients, nutrient availability, and immune effectors^5, 6^. Despite advances in high-throughput sequencing technologies, understanding of the spatial organization of complex microbial communities is still limited^7^. Development of highly multiplexed methods for system-level, spatially-resolved profiling of microbial communities is crucial to elucidate principles governing the assembly and functions of microbiomes, as well as their interactions with the environment and hosts^8-11^.

Fluorescence *in situ* hybridization (FISH) with probes targeting ribosomal RNA (rRNA) has been widely used to identify specific microbial taxa and allows for *in situ* spatial analysis of microbiomes at single cell resolution^12-16^. One challenge of this spatial profiling is the huge phylogenetic and functional diversity of free-living and host-associated microbial communities. Hundreds to thousands of microbial species reside in soil, plant rhizospheres^17^, and mammalian guts^18^. *In situ* profiling of meta-transcriptomes is even more challenging, as the estimated complexity of metagenomes (e.g., over 20 million genes in the human gut microbiome^19^) far exceeds the complexity of host genomes. Multiple methods have been developed to increase multiplexity in spatial mapping of microbiomes^20-25^. Combinatorial labeling and spectral imaging in CLASI-FISH allowed the number of imaged species to exceed the number of fluorophores^21, 22^. The two-step hybridization scheme and advanced spectral unmixing in HiPR-FISH further increased multiplexity^25^. However, the multiplexity of currently available imaging methods for microbiome samples is still inherently limited by the number of fluorophores, i.e., up to 2^F^-1 targets with F fluorophores (Supplementary Table 1). In contrast, technology in spatial genomics of mammalian cells have exploited multiple rounds of hybridization and imaging^26-30^. Methods based on sequential FISH can significantly increase multiplexity and are very much needed for spatial metagenomics of microbial communities.

Another challenge of applying FISH methods to profile complex microbial communities is the accuracy of target identification. The accuracy of taxonomic identification with FISH depends on the specificity of probe targeting, yet rRNA sequences of species closely related phylogenetically are highly similar. In a diverse microbial community, probes of nonspecific binding cannot be ignored; therefore, it is difficult to design perfectly selective probes at the species or sub-species level^11, 31^, and to achieve highly accurate target identification with multiplexed FISH methods. Incorporating error-correction strategies is expected to improve accuracy^29, 30^, but has not been studied in the context of microbiome imaging.

Here we introduce sequential error-robust fluorescence *in situ* hybridization (SEER-FISH), a highly multiplexed and accurate microbiome imaging approach allowing spatial mapping of microbial communities at single cell resolution. We developed a protocol that allows for multiple rounds of FISH imaging of microbial samples. The exponential combination of fluorophore numbers (F) and hybridization rounds (N) leads to an unparalleled increase in multiplexity (F^N^) for labeling microbiome samples. By incorporating error-robust encoding schemes, we show that SEER-FISH could tolerate probe non-specificity to achieve high precision and recall in taxonomic identification of microbial communities. Finally, as a proof-of-concept demonstration, we applied SEER-FISH in imaging *Arabidopsis thaliana* roots to unravel the micron-scale biogeography of microbial communities colonizing the rhizoplane.

## Results

### Superior multiplexity of SEER-FISH in spatial profiling of microbiome

SEER-FISH provides a scalable coding capacity of F^N^ (F-color, N-round) by encoding each target taxon with a unique barcode through N rounds of imaging (Fig. 1A). To sequentially label the target taxon with an N-bit barcode, we developed a protocol that allowed for iterative labeling of microbial rRNAs with rapid probe hybridization and dissociation (Supplementary Fig. 1, see Methods for details). In each round, probes of F colors were hybridized to targeted rRNAs, and the sample was imaged and then treated with dissociation buffer to remove the hybridized probes^28, 32^. After experiments were completed, the images were aligned to eliminate the shift in position during multiple rounds of imaging. The boundaries of bacterial cells were segmented using the watershed algorithm, and the fluorescence intensity of bacterial cells in each round of imaging was determined to identify their corresponding barcodes (Supplementary Fig. 2).

**Figure 1.**
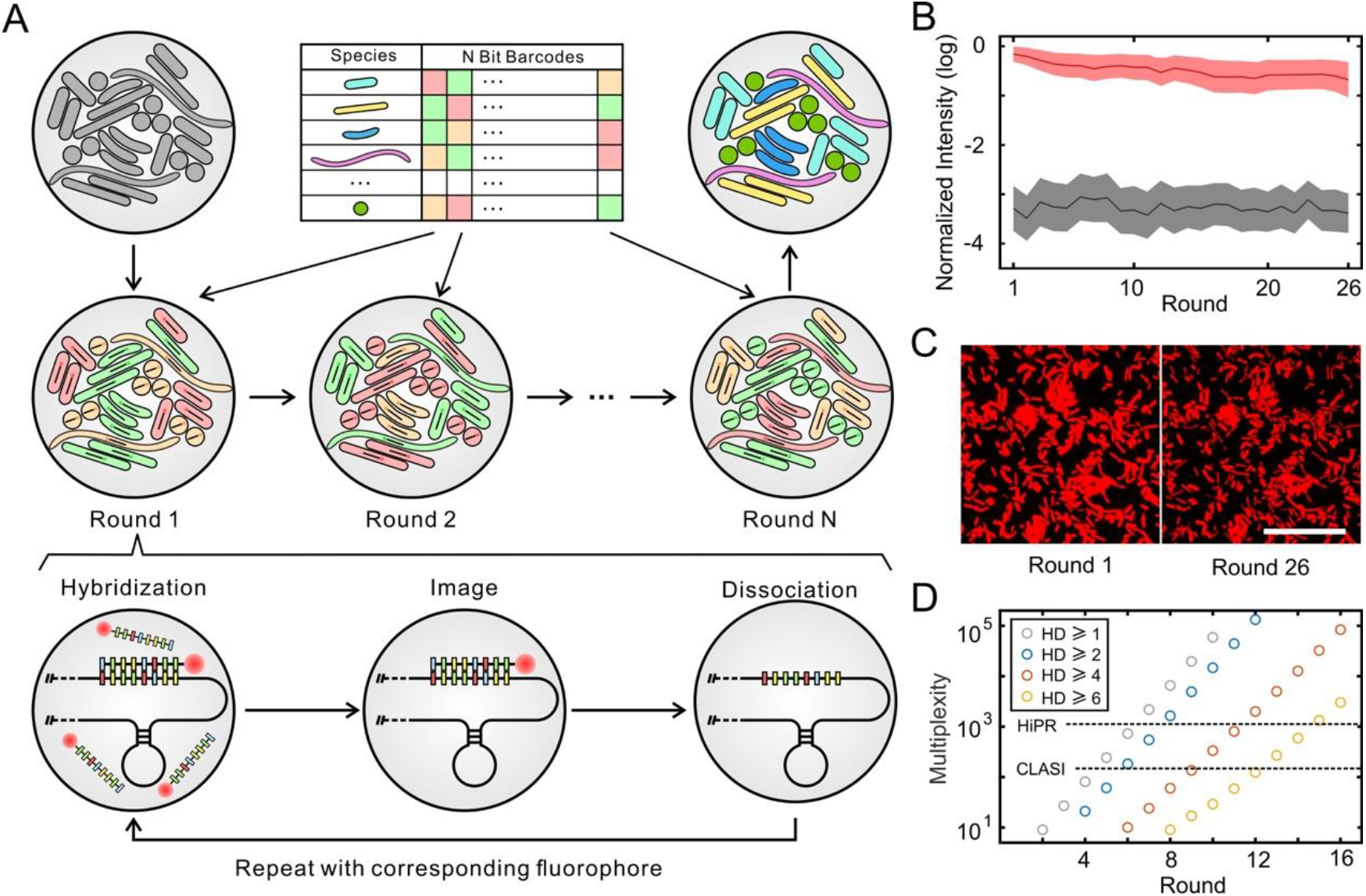
SEER-FISH allows superior multiplexity in spatial profiling of microbiomes. **A)** Design of SEER-FISH. Each bacterial taxon is encoded by an F-color N-bit barcode. The spatial distribution of the microbial community can be obtained through R rounds of FISH. Each round of SEER-FISH includes probe hybridization, imaging, and probe dissociation (Methods and Supplementary Fig. 1A). **B)** Fluorescence intensity over 26 rounds of SEER-FISH. Lines indicate the mean fluorescence intensity (log-transformed and normalized by the maximum pixel value of CCD) of bacterial cells (n=2,257) after hybridization (red line) and dissociation (black line), respectively. The shadow of each line indicates the standard deviation. **C)** Fluorescence intensities at the 1^st^ and 26^th^ rounds of imaging. Scale bar, 25 μm. **D)** The multiplexity of SEER-FISH increases exponentially with the number of rounds and exceeds the capacity of multi-color combinational labeling methods, e.g. CLASI-FISH (∼100) and HiPR-FISH (∼1,000). The colors of the circles indicate the minimal Hamming distance (HD) between barcodes. All codebooks are generated with three colors (F=3).

To evaluate the feasibility of our experimental protocol, we performed multiple rounds of FISH imaging on a mixture of bacterial species (SynCom12, see Methods) with the universal probe EUB338 (Fig. 1B and C). The hybridized probes were efficiently removed by dissociation buffer, leading to an ∼1000-fold decrease in the fluorescence intensity. The dissociation step in our protocol had little effect on subsequent rounds of hybridization (Fig. 1B). In contrast, re-hybridization of probes after photobleaching was inefficient and did not allow for multiple rounds of imaging (Supplementary Fig. 3). After 26 rounds of probe hybridization and dissociation, we found that the mean fluorescence intensity of bacterial cells was still significantly higher than the background (Fig. 1B). Only a small fraction of bacterial cells (less than 3%) were lost due to the decrease in fluorescence intensity or the shift in their location.

Thus, for the first time as we know of, we developed an efficient method to label microbial rRNAs using sequential hybridization and dissociation of probes. Our protocol supports sequential FISH on microbial samples for more than 25 rounds, and each round of imaging takes only ∼15 min (Supplementary Fig. 1). In contrast, other methods of one-round bacterial FISH often require more than 2 hours^14-16, 21-25^. The coding capacity of SEER-FISH, similar to sequential FISH studies in the context of single cell transcriptomics^26-28^, possesses great scalability through increasing the rounds of imaging (F^N^). The multiplexity of SEER-FISH can easily reach 10^5^ (Fig. 1D), which is much higher than existing methods for imaging microbiome samples, such as CLASI-FISH^22^ (multiplexity ∼10^2^) and HiPR-FISH^25^ (multiplexity ∼10^3^). A detailed comparison of different methods can be found in Supplementary Table 1.

### Error-robust encoding enables high accuracy in taxonomic identification

One challenge for sequential FISH is that detection errors would increase with rounds of hybridization. Analogous to the strategy previously proposed in the context of labeling mRNAs in mammalian cells^29, 30^, we designed an error-robust encoding scheme that used a subset of the F^N^ barcodes with specified minimal Hamming distance (HD) (Fig. 1D and Fig. 2A, Methods). For example, any two barcodes from a codebook with a minimal HD of 4 (HD4) differ by at least 4 bits. Therefore, we can correct 1-bit and some 2-bit detection errors by identifying the observed barcode compared to its nearest valid barcodes (Fig. 2B). Each bacterial taxon is labeled with a specific fluorescence color in each round, e.g., FAM/Cy3/Cy5(F=1/2/3), which is decoded by comparing brightness across different fluorescence channels (Methods).

**Figure 2.**
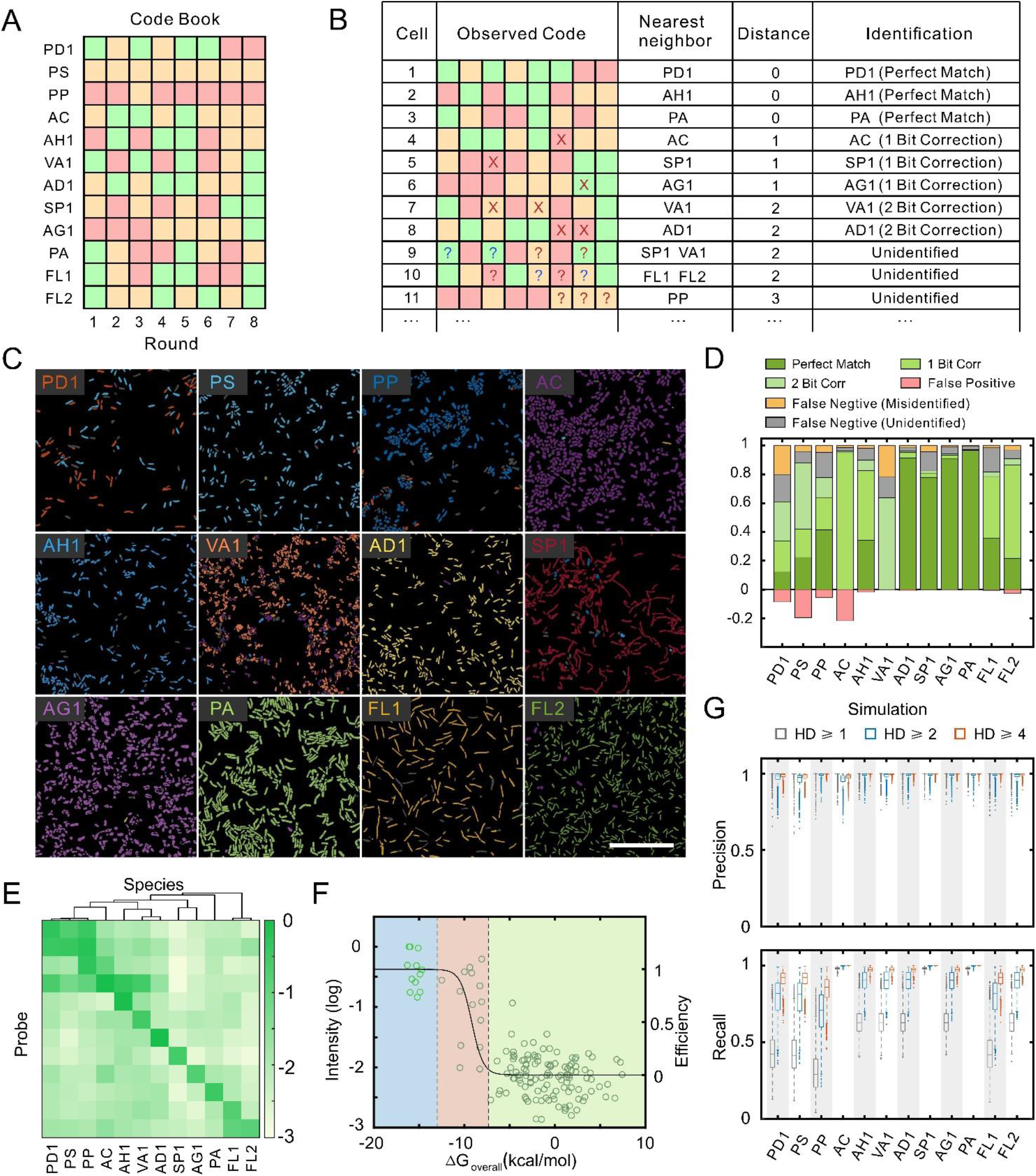
SEER-FISH enables highly accurate taxonomic identification. **A**) The codebook used for the validation experiment on the synthetic community consisting of 12 bacterial species (Supplementary Table 5, R8HD4 codebook, the barcodes for a specific set of strains (S): R=8, HD≥4, S=12). **B)** Illustration of the decoding scheme for the codebook shown in panel A. Crosses (or question marks) indicate errors that can (or cannot) be corrected by mapping to the nearest neighbor in the codebook. **C)** Identification of bacterial species grown in pure culture by eight rounds of imaging (R8HD4 codebook). The pseudocolor of each bacterial species is indicated by its acronym. Scale bar, 25 μm. **D)** Quantification of results in panel C. For each species, cells correctly identified (including perfect match, 1-bit correction, and 2-bit correction) are true positives (Green);cells incorrectly identified as the other 11 species are marked as misidentified (Orange); cells that cannot be classified to any of the 12 species are marked as unidentified (Gray). Cells of the other 11 species incorrectly identified as the corresponding species are false positives (Red). The ratios of true positives, misidentified cells, and unidentified cells sum to 1. Ratios are normalized by the total cell number of each species. **E)** Analysis of probe specificity. The measured fluorescence intensity of bacterial cells (pure culture, average of ∼1,000 cells) hybridized with probes designed to target individual species. The species are clustered by the phylogenetic distance between full 16S sequences (Minimum-Evolution Tree, MEGA-X v10.1.8). Probes (y-axis) follow the same order as the targeted species (x-axis). **F)** The relationship between the measured fluorescence intensity and the change in free energy (Δ*G*) of each probe-species pair (Methods). The light and dark green circles indicate the measured fluorescence intensity of diagonal and off-diagonal probe-species hybridization shown in panel E, respectively. The black line indicates the predicted hybridization efficiency *E* (Methods). The colored regions indicate specific binding (Δ*G* < -13.0 kcal/mol, blue), non-specific binding (−7.9 kcal/mol > Δ*G* >-13.0 kcal/mol, red), and background (Δ*G* >-7.9 kcal/mol, green) (Supplementary Fig. 6). **G)** Simulations show that both precision and recall of taxonomic identification are improved by error-robust encoding. The colored boxplot indicates the predicted distribution of precision and recall of SEER-FISH with 5,000 randomly generated codebooks (F=3, R=8, S=12) with different minimal HD (HD≥1, 2, 4). The height of the box indicates the first and third quartiles.

To evaluate the feasibility of error-robust encoding schemes, we performed SEER-FISH on pure cultures of 12 bacterial species (Supplementary Table 2) with a set of R8HD4 barcodes (rounds=8, minimal HD=4, Fig. 2A). The design and selection of probes specifically targeting 16S or 23S rRNA of the corresponding bacterial species is based on stringent criteria that take into account sequence mismatch to non-target taxa^33^ and predicted hybridization efficiency to the targeted taxa^34^ (Supplementary Fig. 4). Each bacterial species was separately coated onto a coverslip, hybridized with probes according to the codebook and imaged for eight sequential rounds. Finally, bacterial cells were identified by decoding their barcodes and compared with ground truth (Fig. 2C and D, Supplementary Fig. 5). We found that SEER-FISH had excellent precision (median=0.98, ranging from 0.78 to 0.99) and recall (median=0.89, ranging from 0.61 to 0.97) in bacterial identification (Methods, Supplementary Fig. 5C), as most of the cells were correctly identified. In particular, we found that recall was significantly improved via error correction (27% via 1-bit correction, and 14% via 2-bit correction); otherwise, these observed barcodes would be unidentified because of detection errors (Fig. 2D).

Despite the stringent criteria used in probe design, due to the sequence similarity between closely related bacterial species and complex effects of sequence mismatch on hybridization efficiency, non-specific binding in bacterial rRNA FISH is difficult to be completely eliminated^31^. To investigate how probe non-specificity influenced the performance of SEER-FISH, we systematically profiled the specificity matrix (12 probes vs. 12 bacterial species) using conventional one-step hybridization FISH (Fig. 2E, Supplementary Fig. 6). We found several cases of non-specific hybridization (i.e., off-diagonal fluorescence signal in the specificity matrix), especially for phylogenetically related species. Non-specific binding of probes caused low precision for species PS and AC (due to false positives) and low recall for species PD1 and VA1 (due to false negatives) (Supplementary Fig. 5C, see species list in Supplementary Table 2). Furthermore, we found that the measured fluorescence intensity was in good agreement with the predicted hybridization efficiency^34^ (more negative Δ*G* means better probe hybridization) (Fig. 2F).

To systematically evaluate the importance of error-robust encoding for sequential FISH with probe non-specificity, we performed simulations for randomly generated codebooks with variable minimal HD (Methods). In the simulated experiments, we classified the probe-species pairs into three groups based on binding free energy Δ*G*, namely specific binding, non-specific binding, and background. In particular, non-specific binding of probes (−13.0 < Δ*G* < -7.3 kcal/mol) could lead to detection errors. At an intermediate level of non-specificity (Fig. 2G, Methods), simulation results showed that error-robust encoding (i.e., increased minimal HD) enabled overall improvements in precision and recall of bacterial identification. Qualitatively similar results were observed in simulations at different levels of non-specificity (Supplementary Fig. 7). In summary, our experimental and computational results showed that sequential FISH with error-robust encoding could tolerate probe non-specificity to provide accurate and sensitive identification of bacterial taxa.

### Profiling the composition of microbial communities by SEER-FISH

To evaluate the reproducibility of taxonomic profiling in microbial communities using SEER-FISH, we performed benchmarking experiments on synthetic communities consisting of 12 species (SynCom12, SynCom12_unequal) and 30 species (SynCom30) (Supplementary Table 2, Methods). Using the R8HD4 codebook (rounds = 8, minimal HD = 4, the barcodes for a specific set of strains (S) = 12, Supplementary Table 5), all 12 species were successfully identified in SynCom12 (Fig. 3A). We found close agreement in the estimated taxonomic composition based on SEER-FISH across different Fields of View and experimental replicates (Pearson correlation R≥0.9) (Fig. 3B). Moreover, we altered the relative abundances of four species in the synthetic community (SynCom12_unequal), where the proportions of FL1 and AD1 were increased to 15.7% and the proportions of AC and PA were decreased to 1%. We found that SEER-FISH accurately quantified changes in the community composition (Fig. 3C).

**Figure 3.**
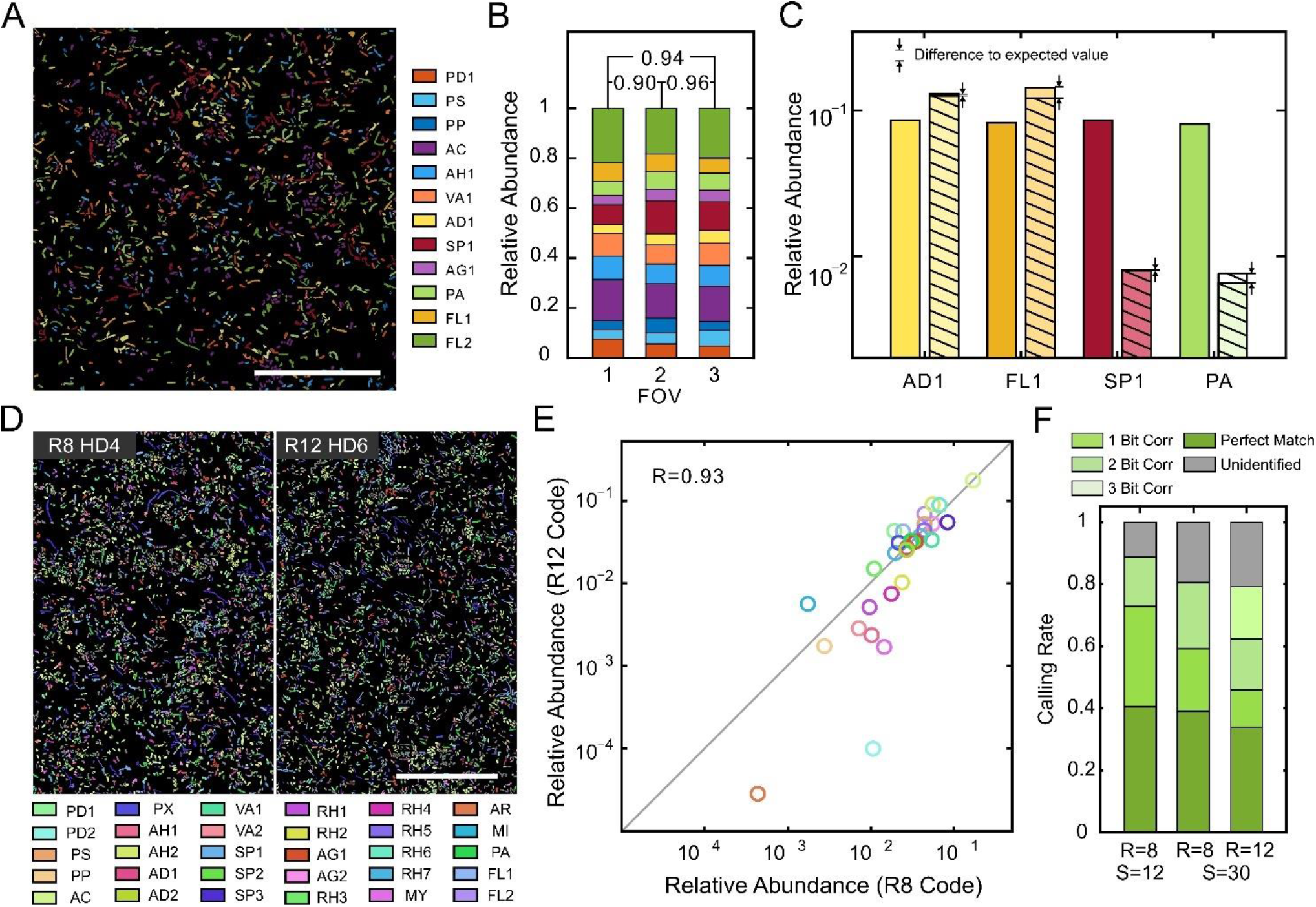
SEER-FISH gives robust estimates of the composition of complex microbial communities. **A)** Representative image, profiling of SynCom12 based on R8HD4 codebook. Scale bar, 50 μm. **B)** Quantification of 12 species relative abundance in SynCom12 in three independent imaging experiments (n=15818, 33365 and 24503 cells, respectively). Pearson correlation between different Fields of View (2 and 3) and between experimental replicates (1 and 2, 1 and 3) are indicated. **C)** Bars of the same color indicate the relative abundance of a given species in SynCom12 (left) and SynCom12_unequal (right) quantified by SEER-FISH. The expected relative abundance after adjustment in SynCom12_unequal is labeled by the stripes. **D)** Representative images, profiling of SynCom30 based on two different codebooks (R8HD4, n=93596 cells vs. R12HD6, n=101084 cells). Scale bar, 50 μm. **E)** Correlation between the relative abundance profiles estimated by two imaging experiments using codebook 1 (R=8, HD≥4, S=30) and codebook 2 (R=12, HD≥6, S=30). **F)** The ratios of identified and unidentified cells. Left bar: SynCom12 (panel A); middle and right bars: SynCom30 (panel D).

Furthermore, we profiled a more complex microbial community (SynCom30) to evaluate the performance of SEER-FISH under different encoding schemes. We chose two codebooks, R8HD4 (rounds = 8, minimal HD = 4, S = 30) and R12HD6 (rounds = 12, minimal HD = 6, S = 30) with high F1 scores (i.e., the harmonic mean of precision and recall) predicted by simulations (Supplementary Fig. 8, Supplementary Table 5). All 30 species were successfully identified by SEER-FISH in both R8HD4 and R12HD6 codebooks (Fig. 3D), and the estimated compositional profiles were highly correlated (Fig. 3E, Pearson correlation R=0.93). In both codebooks, SEER-FISH identified ∼80% of the bacterial cells in the community (Fig. 3F). For R12HD6 codebooks, the increase in imaging rounds led to more detection errors, yet its minimal HD allowed for error correction up to 3 bits and a higher F1 score than R8HD4 codebooks (Supplementary Fig. 8D). Similar to the simulation results for 12 species (Supplementary Fig. 7), we found that error-robust encoding led to enhanced accuracy in taxonomic identification for the 30-species microbial community (Supplementary Fig. 8). Overall, we found that SEER-FISH can be used to quantify the composition of complex microbial communities and that such profiling is highly reproducible.

### Spatial mapping of microbial biogeography on *Arabidopsis* rhizoplane

Spatial distribution of the root microbiome influences plant physiology and development^6, 35-37^. FISH-based methods have been used to characterize root-inhabiting bacteria on the rhizoplane, yet with limited taxonomic resolution^15^. Here we implemented SEER-FISH on *Arabidopsis* roots to map the biogeography of microbial communities colonized on the rhizoplane and to demonstrate the utility of our methods in host-associated microbiome samples.

Axenically grown *Arabidopsis* plants were inoculated with a synthetic community consisting of 12 bacterial species (including ten species isolated from *Arabidopsis* roots^38^ and two *Pseudomonas* strains^39^) and co-cultured for 7 days under hydroponic conditions (Fig. 4A, Methods). After fixation of samples, we imaged the region adjacent to the root tip to quantify the number of bacterial cells and determine the community composition in space (Fig. 4B-E). Over the entire region shown in Fig. 4B (∼1 mm from the root tip), 2,344 bacterial cells (11 out of 12 species) were successfully identified, while 295 cells were unidentified. Only a small fraction of bacterial cells (∼10%) were lost after eight rounds of imaging, indicating that multiple rounds of probe hybridization and dissociation were feasible for tissue samples. There was close agreement between the community composition estimated by SEER-FISH and by 16S rRNA amplicon sequencing of root samples (Pearson correlation R=0.84) (Supplementary Fig. 9A).

**Figure 4.**
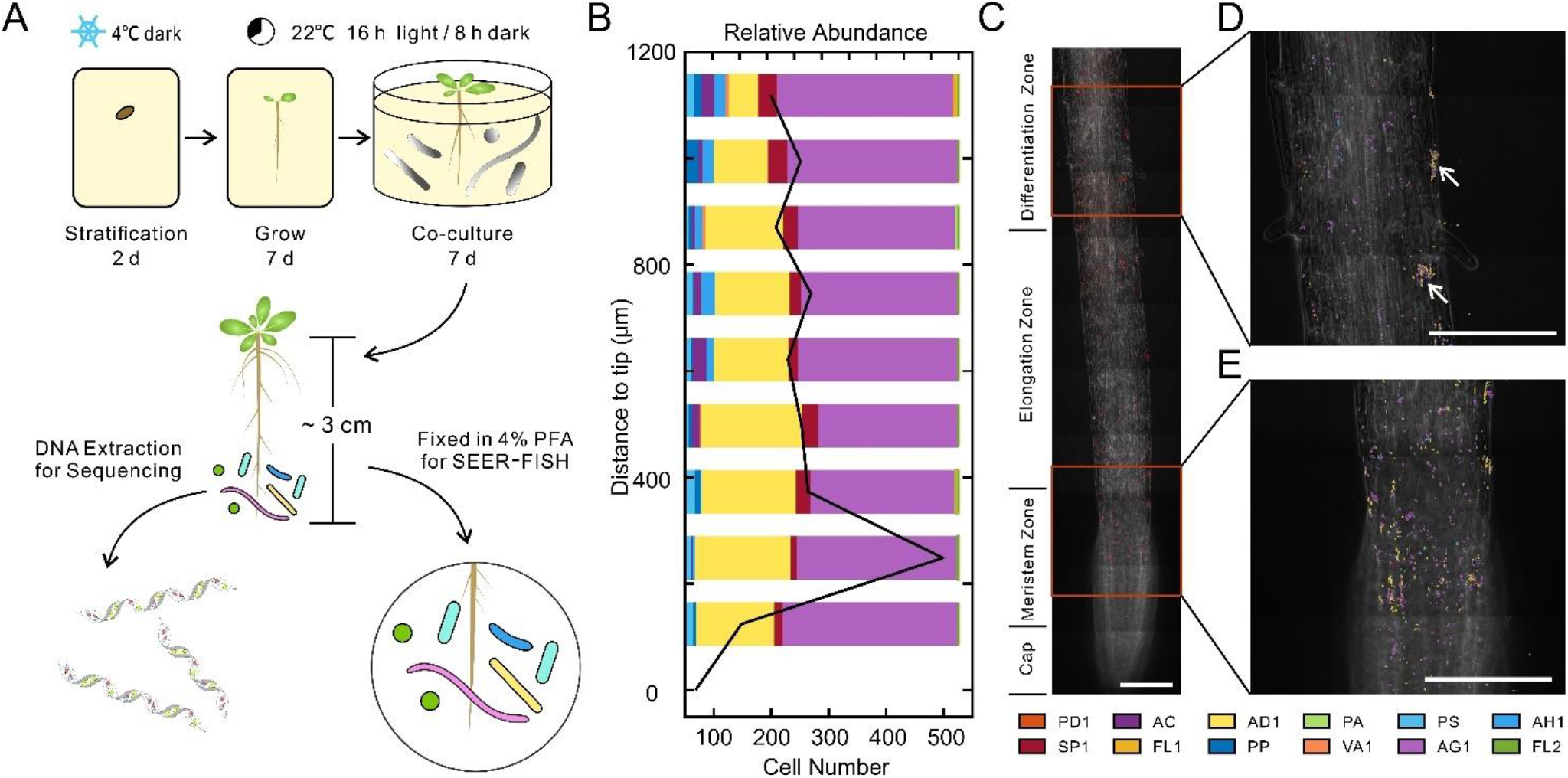
Spatial profiling of microbial communities colonized on *Arabidopsis* roots. **A)** The protocol of synthetic microbial community colonization on *Arabidopsis* roots. *Arabidopsis* seeds were germinated on an MS plate and then colonized by a synthetic community of 12 bacterial species for 7 days under hydroponic conditions (Methods). **B)** Relative abundance of each species (bars) and the number of cells (black line) by imaging along the root of *Arabidopsis*. Each bar indicates the relative abundance in an FOV of 125 μm (width)×250 μm (length). **C)** Representative images illustrate the spatial distribution near the root tip (∼1 mm from the tip) labeled by the universal probe EUB338. Scale bar, 100 μm. **D) and E)** Bacterial colonization marked by the pseudocolor of each species at different regions along the root. Species AG1 and AD1 are found in clusters (white arrows). Scale bar, 100 μm.

In contrast to sequencing, SEER-FISH allowed us to map spatial patterns of the microbiome along plant roots at single cell resolution. We found that the Shannon diversity of the root-inhabiting bacterial communities increased further from the root tip (Supplementary Fig. 9B). The two most abundant species, AD1 (*Acidovorax sp*.) and AG1 (*Agrobacterium sp*.), accounted for ∼80% of the community composition, and were found to aggregate in clusters (Fig. 4D). Moreover, for two regions in the differentiation zone (3-6 mm from root tip, Supplementary Fig. 9 C-F), our imaging revealed accumulation of particular microbial species in root hairs. The differential distribution of microbial communities on the *Arabidopsis* rhizoplane may be linked to localized immune responses or secreted exudates of plant roots^40, 41^. Taken together, our observations highlight the need for comprehensive spatial analysis of microbial colonization on root surfaces.

## Discussion

In this study, we demonstrate that SEER-FISH is a highly multiplexed and accurate imaging technique for investigating the spatial organization of microbiomes. We developed an iterative hybridization and imaging method for microbiome samples. Using error-robust encoding schemes, we show that SEER-FISH provides accurate spatial profiling of microbial communities. While we imaged plant roots as a proof-of-principle demonstration, our methods can be readily applied to image microbiome samples collected from different environments, such as animal guts.

The unparalleled multiplexity of sequential labeling in SEER-FISH combined with the two-step labeling probe design of HiPR-FISH can be used in future studies to take advantage of these complementary approaches (Supplementary Table 1), reducing probe costs and pushing spatial mapping of microbiomes to new frontiers. One exciting prospect is to profile meta-transcriptomes *in situ*, as simultaneous labeling of mRNA and rRNA is feasible for single bacterial strains^13^. The labeling strategy of HiPR-FISH requires a high abundance and uniform distribution of microbial RNA, but this is not necessarily required for SEER-FISH. In addition, the complexity of meta-transcriptomes (>10^7^ genes in human gut microbiomes) require an increase in multiplexity, which can be achieved by sequential labeling. The multiplexity of SEER-FISH can be readily extended by increasing the number of fluorophores and the rounds of imaging. While only three fluorophores were used in this study, our method can easily incorporate more colors and spectral imaging^21, 25^.

By incorporating error-correction strategies in SEER-FISH, we show that the precision and recall of taxonomic identification can be improved, particularly in scenarios where non-specific hybridization is unavoidable. In mRNA labeling, probe specificity is not a major concern and detection errors are dominated by 1→0 errors^29^. In contrast, detection errors in bacterial rRNA FISH are mainly caused by non-specific (i.e., off-target) labeling of phylogenetically related rRNA sequences. In our study, we chose to exclude the non-fluorescent code in the codebook (i.e., color code=0) to minimize detection errors caused by non-specific labeling. Codebooks can be further optimized to account for non-specificity of probes. If the probe specificity matrix (Fig. 2E) of a system has been measured, we can estimate the probability of detection errors and use this information to design better codebooks. Other experimental modifications to reduce non-specific hybridization, such as increasing hybridization stringency^14^, adding competitor probes for off-target taxa^25^ or dual probes with overlapping specificity^31^, can also be used to improve accuracy and are readily compatible with SEER-FISH.

Finally, we envision that the integration of SEER-FISH with other spatially resolved technologies will have broad impacts on microbiology/microbiome research. For example, SEER-FISH can be combined with expansion microscopy to profile the transcriptome of single bacterial cells^42^. Together with mass spectrometry imaging^43^ or multiplexed protein maps^44^, SEER-FISH can unravel the functions of complex microbial communities in space and their interactions with the host at the molecular level.

## Methods

### Bacterial culture

A full list of bacterial strains used in this study is included in Supplementary Table 2. Strains isolated from *Arabidopsis* root microbiota were kindly provided by Professor Paul Schulze-Lefert at the Max Planck Institute for Plant Breeding Research. *Pseudomonas* strains PP (WCS358) and PS (WCS417) were kindly provided by Professor Corné Pieterse at Utrecht University. All strains were cultured in ½ Tryptic Soy Broth (TSB) medium (HuanKai Microbial, 024051) (28°C, shaken at 200 rpm under a normal aerobic atmosphere) and harvested at ∼0.8 OD_600_ (BioTek, Synergy H1 Hybrid Multi-Mode Reader). Bacterial cells were centrifuged at 5,000 ’g for 5 minutes, resuspended and washed with 1× PBS (Boster, AR0030). After that, bacterial cells were centrifuged, resuspended, and fixed with 4% paraformaldehyde (DF0135-2; Leagene) in 1× PBS at 4°C for 3 hours. Fixed bacterial cells were washed 2-3 times in PBS, resuspended in 1× PBS, and then stored in 50% ethanol at -20°C before imaging. For the synthetic community of 12 bacterial strains, two different compositions were created: SynCom12 (mixed with equal OD_600_): 8.3% for each strain. 2) Syncom12_unequal (mixed with unequal OD_600_): the proportions of *Flavobacterium sp*. (FL1) and *Acidovorax sp*. (AD1) were 15.7%; the proportions of *Acinetobacter sp*. (AC) and *Paenibacillus sp*. (PA) were 1%; and the proportions of other strains were 8.3%. For the synthetic community of 30 bacteria strains (SynCom30), all strains were mixed at equal OD_600_.

### Probe design

All probes used in this study are listed in Supplementary Table 3. Oligonucleotide probes were conjugated with three different types of fluorophores at the 5’ terminus: FAM, Cy3, and Cy5 (ordered from General Biol). Oligonucleotide probes were designed to target 16S rRNA or 23S rRNA using a custom pipeline (Supplementary Fig. 4). rRNA and mRNA sequences of the bacterial strains used in this study were extracted from whole genome sequencing data using Prokka^45^, built into a local database and imported to the ARB program (www.arb-home.de). The ‘Probe Design’ function of ARB was used (parameter settings: 18-21 nucleotides, 45-60% GC content)^33^. Probes with fewer than three mismatches to non-target sequences were excluded. The change in free energy (Δ*G*) for probe-target binding was calculated by mathFISH^34^ and probes with Δ*G*<-13.0 kcal/mol were chosen as candidate probes.

The hybridization efficiency *E* is predicted as (Fig. 2F):

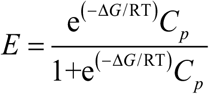

where *C*_*p*_ is the probe concentration of, *R* is the molar gas constant and *T* is the temperature.

### Codebook generation

The codebook is named according to the number of imaging rounds (R), colors (F), and the minimal Hamming distance (HD). All codebooks used in this study contain three colors (F=3); the color code equals 1, 2, and 3 for FAM, Cy3 and Cy5 fluorophores, respectively. First, all *F*^*R*^ barcodes for R rounds of hybridization were generated. Then, the error-robust codebook was generated by repeated removal of barcodes whose distance to the seed barcode was less than the specified minimal HD. Each barcode in the codebook was taken as the seed code until the distance between any two barcodes in the codebook was equal to or larger than the specified minimal HD. The barcodes for a specific set of strains (S) were randomly drawn from the codebook. The codebooks used in the experiments of this study are included in Supplementary Table 5.

### Coverslip functionalization

Coverslips (40-mm, #1.5; Bioptechs) were immersed in potassium dichromate concentrated sulfuric acid cleaning solution for 2 hours, washed with water and then rinsed with distilled water more than three times, soaked in 95% ethanol for 12 hours, and then air-dried. 50X Anti-Slice Escaping Agentia APES (Sangon Biotech, E676003) was diluted 1:50 with acetone and the prepared working solution was used immediately. The coverslips were dipped into the freshly prepared working solution for 20-30s, then removed, followed by a pause before they were washed three times with distilled water to rinse unbound APES. Adhesive processed coverslips were put in a dust-free environment and kept dry.

### Multi-round FISH imaging of bacterial communities

An adhesive coverslip coated with fixed cells was assembled into a Bioptechs FCS2 flow chamber with temperature control (Supplementary Fig. 1B). Fluidics was controlled via a peristaltic pump (LongerPump, BT100-2J) and set at a constant flow velocity of 500 µl/min (10 rpm). Multiple rounds of probe hybridization, imaging, and probe dissociation were performed as follows (Supplementary Fig. 1A):

#### 1) Probe hybridization and washing

1 mL of hybridization buffer (0.9 M NaCl, 0.02 M Tris-HCl (pH 7.6), 0.01% SDS, and 20% formamide (Aladdin, F103362)) with probes was flowed through the sample for 2 min and then incubated for 3 min at 46°C. Probes used in each round were determined by the codebook. Samples were washed by washing buffer (0.215 M NaCl, 0.02 M Tris-HCl (pH 7.6), 0.01% SDS, 5 µM EDTA) for 2 min at 46°C to eliminate residual and nonspecific binding of the probes.

#### 2) Imaging

Images were acquired by a confocal laser scanning microscope (Nikon, A1) with a Plan Apo λ 100 x oil objective lens (Nikon 1.5 NA). Multiple (∼4 to 36) fields of view (125 µm by 125 µm) were collected by sequential excitation with laser lines 488 nm, 561 nm, and 640 nm. Phase-contrast images were acquired by a transmitted detector using a 640 nm laser. The constant focus during imaging was achieved by Nikon PFS autofocus. The image acquisition settings are listed in Supplementary Table 4.

#### 3) Probe dissociation

dissociation buffer (70% formamide, 0.02 M Tris-HCl (pH 7.6), 0.01% SDS, 5 µM EDTA) was flowed through the samples at 46°C for 2 min to strip off hybridized probes.

### Image analysis

First, phase contrast images of each round were used for alignment. The images were aligned to the position with maximum cross correlation by custom MATLAB scripts to eliminate the position shift during multiple rounds of imaging. Then, the phase contrast image (for *in vitro* bacterial communities) or inverted fluorescence image with the universal probe EUB338 (for root-associated bacterial communities) were segmented into binary mask images using an adaptation threshold^46^ followed by the watershed algorithm^47^ (Supplementary Fig. 2). Finally, the fluorescence intensity of each cell was obtained with the mask generated by image segmentation. The color code of each cell in each round was determined by the brightest fluorescence channel in the corresponding round (color codes equal 1, 2, or 3 for FAM, Cy3, or Cy5 channels, respectively). If the fluorescence intensity was not significantly brighter than the background in all fluorescence channels (p>0.05, t-test), the code of the corresponding bacterial cell was marked as 0. A cell was marked as lost if there were more than three zeroes in its barcode.

### Barcode identification

The barcode of each bacterial cell was mapped to the codebook (Supplementary Table 5) to find the nearest neighbor. For barcodes with minimal HD of 2k (k=1, 2,…), if the observed barcode was the same as (perfect match) or had less than n-bit difference (n-bit correction, n<k) from the nearest neighbor, the bacterial cell was successfully identified. If the observed barcode had k-bit difference from the nearest neighbor in the codebook and the nearest neighbor was unique, the cell was also successfully identified (k-bit correction). Otherwise, the cell was labeled as unidentified. For example (Fig. 2B), if there was a 1 bit difference between the observed barcode and the nearest candidate barcode, the corresponding cell was identified by 1-bit correction. If there were 2 bit differences between the observed barcode and the nearest candidate barcode, the error could be corrected only when there were 3 or more bits of difference for other candidate barcodes. The cell was marked as unclassified (unidentified) for other conditions.

### Precision and recall calculation

The precision and recall of each bacterial species are calculated by the following equations:

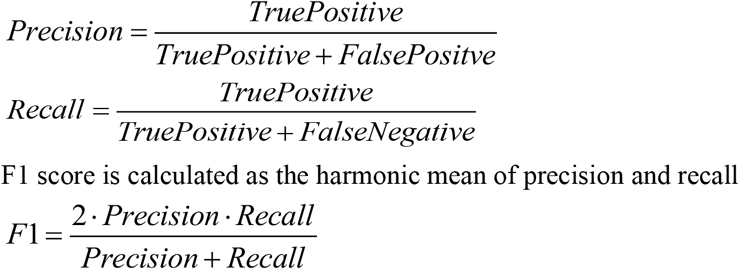

### Imaging of bacterial communities on *Arabidopsis* roots

Surface sterilized *Arabidopsis* seeds of wild-type (Col-0) were sown on 1× Murashige and Skoog (MS) medium (Solarbio, M8520) with 3% sucrose and 0.6% agar. After 2 days of cold-stratification at 4°C under darkness, the plates were then kept in a growth chamber (22°C, 16h light/8h dark, 50% humidity) for 7 days. Twelve species of selected bacteria were pre-cultured with ½ TSB as previously described. Cells were collected by centrifuging and then washed with 1× PBS. The OD_600_ of each strain was determined with the BioTek plate reader. The synthetic bacterial community (OD_600_ = 0.01) containing all strains at equal proportions was inoculated in 1× MS liquid media. Seven-day old seedlings were transferred into the 12-well plate using sterilized tweezers to co-culture with the bacterial community. After 7 days of bacteria-plant co-culture, the roots of seedlings were fixed with 4% paraformaldehyde (PFA) in 1× PBS at 4°C for 3 h. Fixed roots were rinsed and then stored in 50% ethanol at -20°C. Multi-round FISH imaging was carried out as described above. All bacteria were labeled with the universal bacterial probe EUB338 during the first round of imaging. The duration of probe hybridization was extended to 8 min for root samples.

### 16S amplicon sequencing of synthetic bacterial communities

Roots of *Arabidopsis* were harvested, rinsed in 1× PBS buffer and stored at -80°C until further analysis. Each sample of ten frozen roots was treated in liquid nitrogen briefly before manual homogenization using plastic pestles. Genomic DNA was extracted with the DNeasy Plant Mini Kit (QIAGEN GmbH, 69106). The sample was amplified with barcode primers 799F (5′AACMGGATTAGATACCCKG-3′) and 1193R (5′ACGTCATCCCCACCTTCC-3′) against the V5-V7 region of the bacterial 16S rRNA gene. The 25 μl PCR reaction contained 12.5 μL of 2× PrimeSTAR Max Premix (TaKaRa, R045A), 0.4 μM of each primer (Genewiz), and 1 ng genomic DNA. PCR conditions were set as: 1) 98°C for 5 min; 2) 35 cycles of 98 °C for 10s, 50°C for 15s, and 72°C for 15s; 3) elongation at 72°C for 5 min. PCR products were purified using a Gel Extraction Kit (OMEGA, D2500-01) and quantified with Qubit 4 fluorometer (Invitrogen). The amplicon library was sequenced by Illumina MiSeq (500-cycle, V2 kit). Raw 16S rRNA amplicon sequence data were processed by QIIME2^48^. The forward and reverse reads were merged by VSEARCH^49^. Reads were demultiplexed and aligned to a reference set of 16S amplicon sequences to calculate the relative abundance of each taxon using custom scripts.

### Simulations

The intensity of each bacterial cell of species *i* in channel *k* at each round is

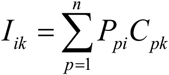

*P*_*pi*_ is the fluorescence intensity when probe *p* is hybridized to species *i. C*_*pk*_ (=1 present, 0 absent) indicates the code status of probe *p* in fluorescence channel *k*. The strength of hybridization was assumed to be the same in different fluorescence channels and classified into three groups (specific binding, non-specific binding, and background), according to the overall Gibbs free energy change (ΔG) when probe *p* hybridized to species *i*. The log-transformed fluorescence intensity (log(*P*_*pi*_)) is drawn from a normal distribution. The mean log-transformed fluorescence intensity is set to -0.3, -0.9 and -2 for specific binding, non-specific binding and background, respectively (Fig. 2G, Supplementary Figure 6C). The standard deviation of log(*P*_*pi*_) was set as 0.3 (Supplementary Figure 6D). For simulations shown in Supplementary Figs. 7 and 8, the mean log-transformed fluorescence intensity for non-specific binding was set to -0.6 for strong non-specificity, -1.2 for weak non-specificity. The fluorescence intensity of each cell in each fluorescence channel was drawn independently during each round according to the codebook, and the barcode was identified as described above.

### Microscopy data availability

All microscopy data will be deposited to Zenodo.

### Code availability

All scripts will be available on Github.

## Acknowledgments

We thank Prof. Yang Bai at the Institute of Genetic and Developmental Biology, Chinese Academy of Sciences for help with strain shipping and cultivation, plant experiments and critical comments on the manuscript. We thank Dr. Ziyu Li for technical advice and assistance with instrumentation. We thank members of the LD lab for insightful discussions. We thank Ke Yu, Yuqian Wu, Yang Bai, Sihong Li at SIAT for their help during the early stage of this project.

## Author contributions

L.D. conceived and supervised the study. L.D., Z.C. and W. Z. designed the experiments. Z. C. set up the experimental protocol and performed the imaging experiments. W.Z. performed data analysis and simulations. L.W. performed the plant experiments. Z.Q. assisted with imaging experiments. J.C. assisted with bioinformatics analysis. F.J. assisted with instrumentation setup. L.D., Z. C. and W. Z. wrote the manuscript. All authors discussed the results and commented on the manuscript.

## Notes

### Competing Interest Statement

The authors have declared no competing interest.

## References

1. Nadell, C.D., Drescher, K. & Foster, K.R. Spatial structure, cooperation and competition in biofilms. Nat Rev Microbiol 14, 589–600 (2016).

2. Donaldson, G.P., Lee, S.M. & Mazmanian, S.K. Gut biogeography of the bacterial microbiota. Nat Rev Microbiol 14, 20–32 (2016).

3. Mark Welch, J.L., Ramirez-Puebla, S.T. & Borisy, G.G. Oral Microbiome Geography: Micron-Scale Habitat and Niche. Cell Host Microbe 28, 160–168 (2020).

4. Richardson, A.E., Kawasaki, A., Condron, L.M., Ryan, P.R. & Gupta, V.V.S.R. in Rhizosphere Biology: Interactions Between Microbes and Plants. (eds. V.V.S.R. Gupta & A.K. Sharma) 109–128 (Springer Singapore, Singapore; 2021).

5. Tropini, C., Earle, K.A., Huang, K.C. & Sonnenburg, J.L. The Gut Microbiome: Connecting Spatial Organization to Function. Cell Host Microbe 21, 433–442 (2017).

6. Bulgarelli, D. et al.. Revealing structure and assembly cues for Arabidopsis root-inhabiting bacterial microbiota. Nature 488, 91–95 (2012).

7. Sheth, R.U. et al.. Spatial metagenomic characterization of microbial biogeography in the gut. Nat Biotechnol 37, 877–883 (2019).

8. Cordero, O.X. & Datta, M.S. Microbial interactions and community assembly at microscales. Curr Opin Microbiol 31, 227–234 (2016).

9. Mark Welch, J.L., Hasegawa, Y., McNulty, N.P., Gordon, J.I. & Borisy, G.G. Spatial organization of a model 15-member human gut microbiota established in gnotobiotic mice. Proc Natl Acad Sci U S A 114, E9105–E9114 (2017).

10. Mark Welch, J.L., Rossetti, B.J., Rieken, C.W., Dewhirst, F.E. & Borisy, G.G. Biogeography of a human oral microbiome at the micron scale. Proc Natl Acad Sci U S A 113, E791–800 (2016).

11. Wilbert, S.A., Mark Welch, J.L. & Borisy, G.G. Spatial Ecology of the Human Tongue Dorsum Microbiome. Cell Rep 30, 4003–4015 e4003 (2020).

12. Prudent, E. & Raoult, D. Fluorescence in situ hybridization, a complementary molecular tool for the clinical diagnosis of infectious diseases by intracellular and fastidious bacteria. FEMS Microbiol Rev 43, 88–107 (2019).

13. Pernthaler, A. & Amann, R. Simultaneous fluorescence in situ hybridization of mRNA and rRNA in environmental bacteria. Appl Environ Microbiol 70, 5426–5433 (2004).

14. Thurnheer, T., Gmur, R. & Guggenheim, B. Multiplex FISH analysis of a six-species bacterial biofilm. J Microbiol Methods 56, 37–47 (2004).

15. Schmidt, H. & Eickhorst, T. Detection and quantification of native microbial populations on soil-grown rice roots by catalyzed reporter deposition-fluorescence in situ hybridization. FEMS Microbiol Ecol 87, 390–402 (2014).

16. Kim, D. et al.. Spatial mapping of polymicrobial communities reveals a precise biogeography associated with human dental caries. Proc Natl Acad Sci U S A 117, 12375–12386 (2020).

17. Thompson, L.R. et al.. A communal catalogue reveals Earth’s multiscale microbial diversity. Nature 551, 457–463 (2017).

18. Moeller, A.H. et al.. Dispersal limitation promotes the diversification of the mammalian gut microbiota. Proc Natl Acad Sci U S A 114, 13768–13773 (2017).

19. Tierney, B.T. et al.. The Landscape of Genetic Content in the Gut and Oral Human Microbiome. Cell Host & Microbe 26, 283-295.e288 (2019).

20. Shi, H., Grodner, B. & De Vlaminck, I. Recent advances in tools to map the microbiome. Current Opinion in Biomedical Engineering (2021).

21. Valm, A.M. et al.. Systems-level analysis of microbial community organization through combinatorial labeling and spectral imaging. Proc Natl Acad Sci U S A 108, 4152–4157 (2011).

22. Valm, A.M., Oldenbourg, R. & Borisy, G.G. Multiplexed Spectral Imaging of 120 Different Fluorescent Labels. PLoS One 11, e0158495 (2016).

23. Schimak, M.P. et al.. MiL-FISH: Multilabeled Oligonucleotides for Fluorescence In Situ Hybridization Improve Visualization of Bacterial Cells. Appl Environ Microbiol 82, 62–70 (2016).

24. Lukumbuzya, M., Schmid, M., Pjevac, P. & Daims, H. A Multicolor Fluorescence in situ Hybridization Approach Using an Extended Set of Fluorophores to Visualize Microorganisms. Front Microbiol 10, 1383 (2019).

25. Shi, H. et al.. Highly multiplexed spatial mapping of microbial communities. Nature (2020).

26. Lubeck, E., Coskun, A.F., Zhiyentayev, T., Ahmad, M. & Cai, L. Single-cell in situ RNA profiling by sequential hybridization. Nat Methods 11, 360–361 (2014).

27. Eng, C.L., Shah, S., Thomassie, J. & Cai, L. Profiling the transcriptome with RNA SPOTs. Nat Methods 14, 1153–1155 (2017).

28. Eng, C.L. et al.. Transcriptome-scale super-resolved imaging in tissues by RNA seqFISH. Nature 568, 235–239 (2019).

29. Chen, K.H., Boettiger, A.N., Moffitt, J.R., Wang, S. & Zhuang, X. RNA imaging. Spatially resolved, highly multiplexed RNA profiling in single cells. Science 348, aaa6090 (2015).

30. Moffitt, J.R. et al.. High-throughput single-cell gene-expression profiling with multiplexed error-robust fluorescence in situ hybridization. Proc Natl Acad Sci U S A 113, 11046–11051 (2016).

31. Wright, E.S., Yilmaz, L.S., Corcoran, A.M., Okten, H.E. & Noguera, D.R. Automated design of probes for rRNA-targeted fluorescence in situ hybridization reveals the advantages of using dual probes for accurate identification. Appl Environ Microbiol 80, 5124–5133 (2014).

32. Yilmaz, L.Ş. & Noguera, D.R. Development of thermodynamic models for simulating probe dissociation profiles in fluorescence in situ hybridization. Biotechnology and Bioengineering 96, 349–363 (2007).

33. Ludwig, W. et al.. ARB: a software environment for sequence data. Nucleic Acids Res 32, 1363– 1371 (2004).

34. Yilmaz, L.S., Parnerkar, S. & Noguera, D.R. mathFISH, a web tool that uses thermodynamics-based mathematical models for in silico evaluation of oligonucleotide probes for fluorescence in situ hybridization. Appl Environ Microbiol 77, 1118–1122 (2011).

35. Trivedi, P., Leach, J.E., Tringe, S.G., Sa, T. & Singh, B.K. Plant-microbiome interactions: from community assembly to plant health. Nat Rev Microbiol 18, 607–621 (2020).

36. Mendes, R., Garbeva, P. & Raaijmakers, J.M. The rhizosphere microbiome: significance of plant beneficial, plant pathogenic, and human pathogenic microorganisms. FEMS Microbiol Rev 37, 634–663 (2013).

37. Schmidt, H. et al.. Recognizing Patterns: Spatial Analysis of Observed Microbial Colonization on Root Surfaces. Frontiers in Environmental Science 6 (2018).

38. Bai, Y. et al.. Functional overlap of the Arabidopsis leaf and root microbiota. Nature 528, 364– 369 (2015).

39. Berendsen, R.L. et al.. Unearthing the genomes of plant-beneficial Pseudomonas model strains WCS358, WCS374 and WCS417. BMC Genomics 16, 539 (2015).

40. Zhou, F. et al.. Co-incidence of Damage and Microbial Patterns Controls Localized Immune Responses in Roots. Cell 180, 440–453 e418 (2020).

41. Massalha, H., Korenblum, E., Malitsky, S., Shapiro, O.H. & Aharoni, A. Live imaging of root– bacteria interactions in a microfluidics setup. Proc Natl Acad Sci U S A 114, 4549 (2017).

42. Chen, F. et al.. Nanoscale imaging of RNA with expansion microscopy. Nat Methods 13, 679– 684 (2016).

43. Geier, B. et al.. Spatial metabolomics of in situ host-microbe interactions at the micrometre scale. Nat Microbiol 5, 498–510 (2020).

44. Gut, G., Herrmann, M.D. & Pelkmans, L. Multiplexed protein maps link subcellular organization to cellular states. Science 361(2018).

## References

45. Seemann, T. Prokka: rapid prokaryotic genome annotation. Bioinformatics 30, 2068–2069 (2014).

46. Bradley, D. & Roth, G. Adaptive Thresholding using the Integral Image. Journal of Graphics Tools 12, 13–21 (2007).

47. Shetty, M. & B, R. in 2018 International Conference on Electrical, Electronics, Communication, Computer, and Optimization Techniques (ICEECCOT) 56-59 (2018).

48. Bolyen, E. et al.. Reproducible, interactive, scalable and extensible microbiome data science using QIIME 2. Nature Biotechnology 37, 852–857 (2019).

49. Rognes, T., Flouri, T., Nichols, B., Quince, C. & Mahe, F. VSEARCH: a versatile open source tool for metagenomics. PeerJ 4, e2584 (2016).

