## Supplementary Figures and Tables for "Spatial profiling of microbial communities by sequential FISH with error-robust encoding"

Supplementary Information

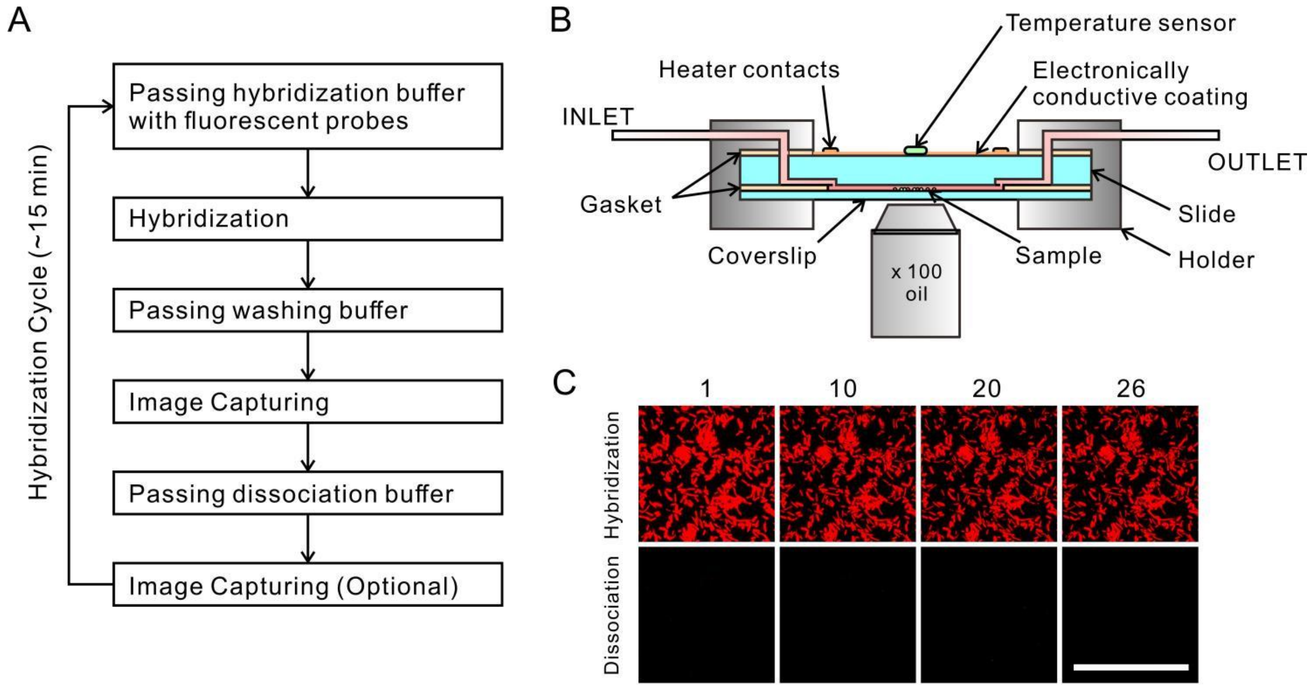


**Figure S1. Sequential FISH imaging of microbial communities. A)** Workflow of SEER-FISH experiments. For each round of hybridization, hybridization buffer with fluorescent probes flow through the sample for 2 min, flow is stopped and sample is incubated for 3 min at 46°C. Then sample is rinsed with washing buffer for 2 min at 46°C to eliminate residual and nonspecific binding of probes. Images are captured right after the washing. After image acquisition, dissociation buffer is flowed through the samples at 46°C for 2 min to strip off hybridized probes. Then dissociation image is captured with the same parameter above for checking the dissociation efficiency. The whole hybridization cycle can be finished in ~15 min and repeated for N rounds. **B)** Schematic diagram of SEER-FISH experimental setup. A flow chamber (Bioptech FCS2) is secured into a stage adapter to interface with a microscope for imaging. Silicone gasket (40 mm round, 0.75 mm thick) with a rectangle cavity internal that separates the micro-aqueduct slide from the coverslip is used to create an optical cavity in the chamber. Laminar flow perfusion that comes into one of the ports (INLET) on one side of the chamber is collected within the optical cavity and then directed out of the chamber on the other side (OUTLET). Uniform temperature across the entire field is maintained by a temperature controller. Flow through this chamber is controlled via an extraneous peristaltic pump. **C)** Representative images of microbes after multiple rounds of hybridization and dissociation (hybridization round 1, 10, 20, and 26). Dissociation images demonstrate the efficient removal of fluorescent probes. Scale bar, 50 μm.

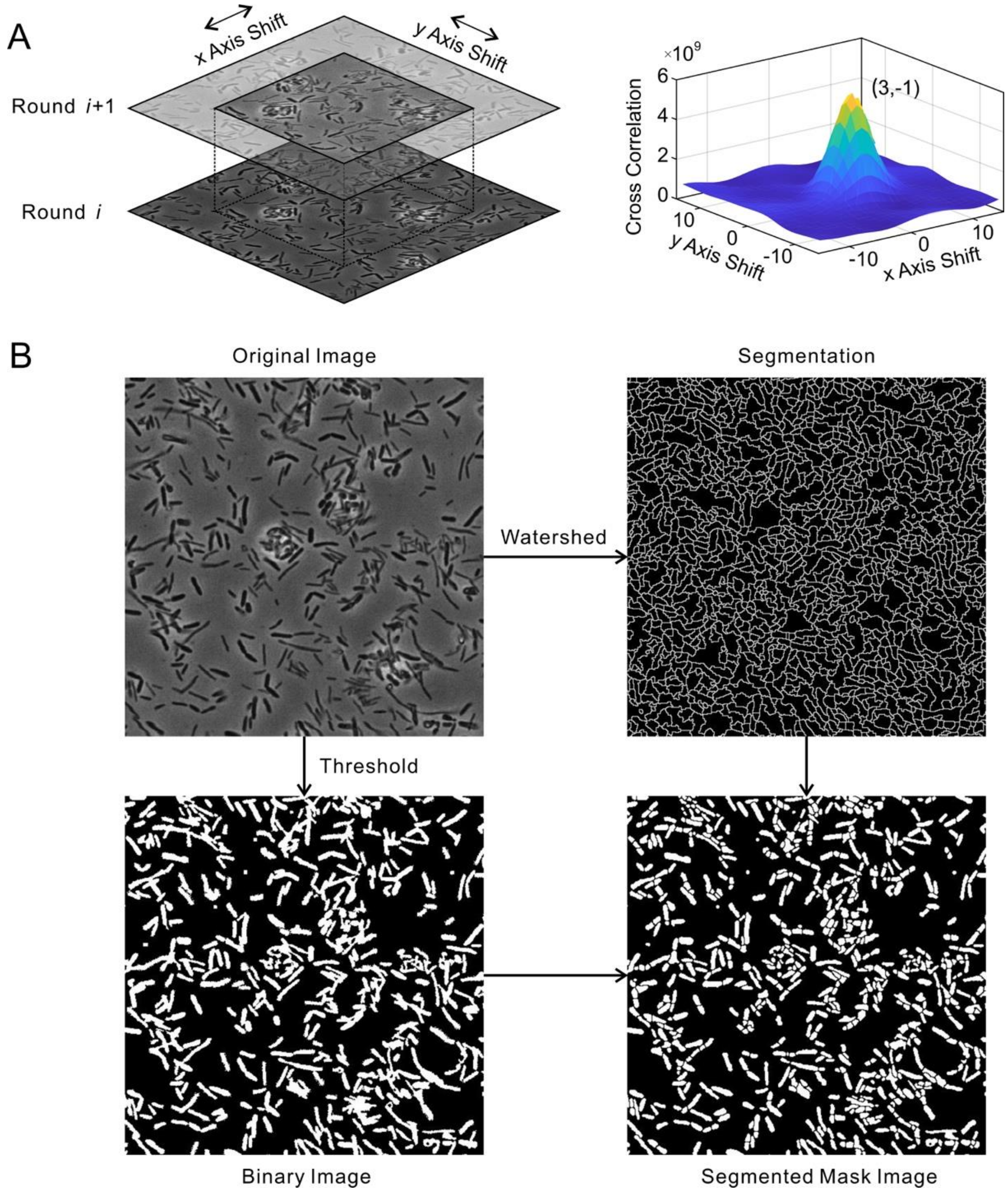


**Figure S2. Workflow for image alignment and segmentation of bacterial cells. A)** The phase contrast images from neighboring rounds are used for alignment. Multiple images were cropped from the center of the round i+1 image with various shift pixels on x and y axis (left panel). The cross correlation between each cropped image and the image of round i is calculated to find cropped image with maximal cross correlation (right panel). The images were aligned by shift the image of round i+1 with the same distance as the cropped image. **B)** Identification of single bacterial cells by segmentation. The binary image is generated with the phase contrast image of the first-round imaging by adaptive threshold. The segmentation is generated by applying the watershed algorithm on the phase contrast image. The final segmented mask image is generated by applying segmentation on the binary image. Images are analyzed by custom MATLAB scripts.

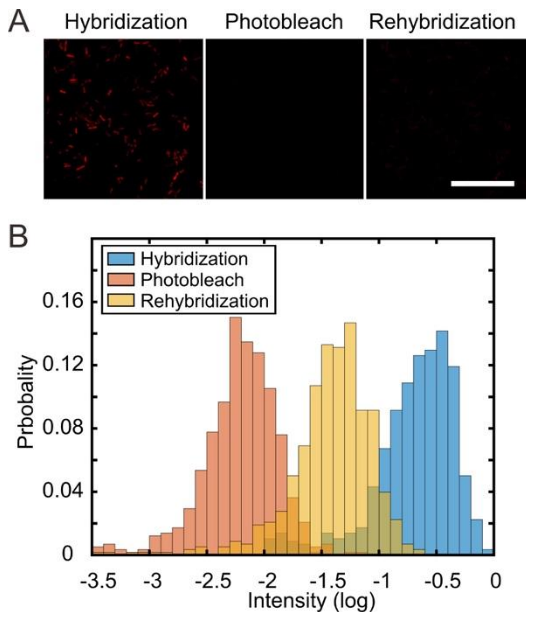


**Figure S3. The rehybridization efficiency is low after photobleaching.** The mixture of 12 bacterial species was labeled by the universal probe (EUB338-Cy5). Fluorescence was eliminated by photobleaching with a high power laser for 2 min. After photobleaching, the sample was rehybridized with same probe. **A)** The fluorescent images after hybridization, photobleaching, and rehybridization. Scale bar, 25 μm. **B)** The distribution of fluorescent intensity of bacterial cells after hybridization, photobleaching, and rehybridization. The mean fluorescent intensity decreased about 30 folds after photobleaching and only recovered to 20% of the original level after rehybridization.

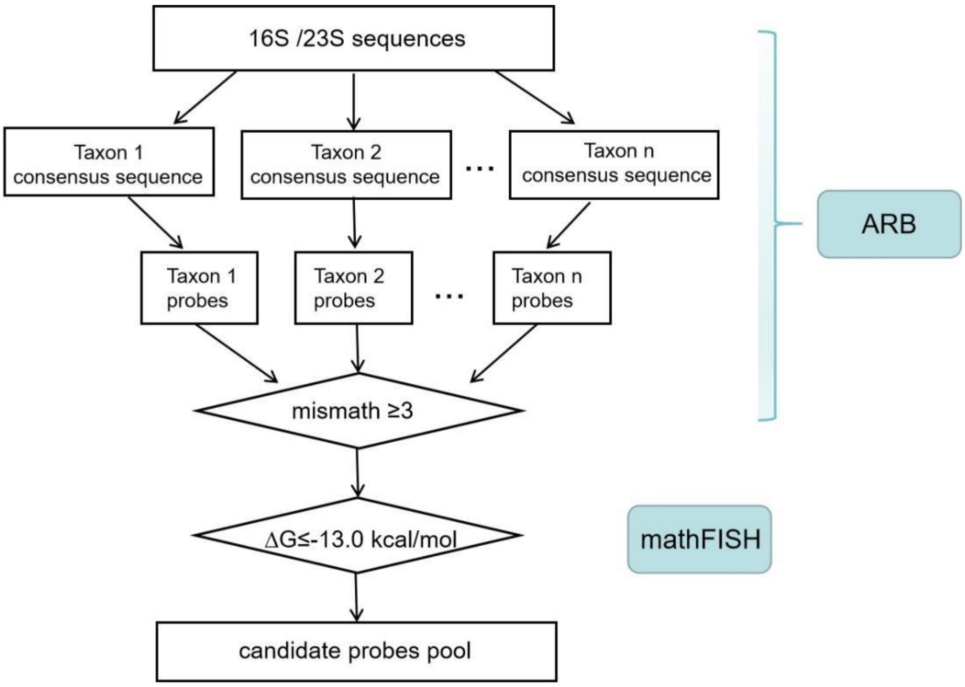


**Figure S4. Design of taxon-specific probes that target bacterial rRNA.** The 16S or 23S rDNA sequences of targeted bacterial taxa are built into a local database and imported to the ARB program for probe design. Probes with at least three central mismatches to all non-target taxa are retained. Furthermore, $\Delta G$ for probes binding to targets is calculated by mathFISH. Probes with a $\Delta G$ value less than -13.0 kcal/mol are chosen as candidate probes.

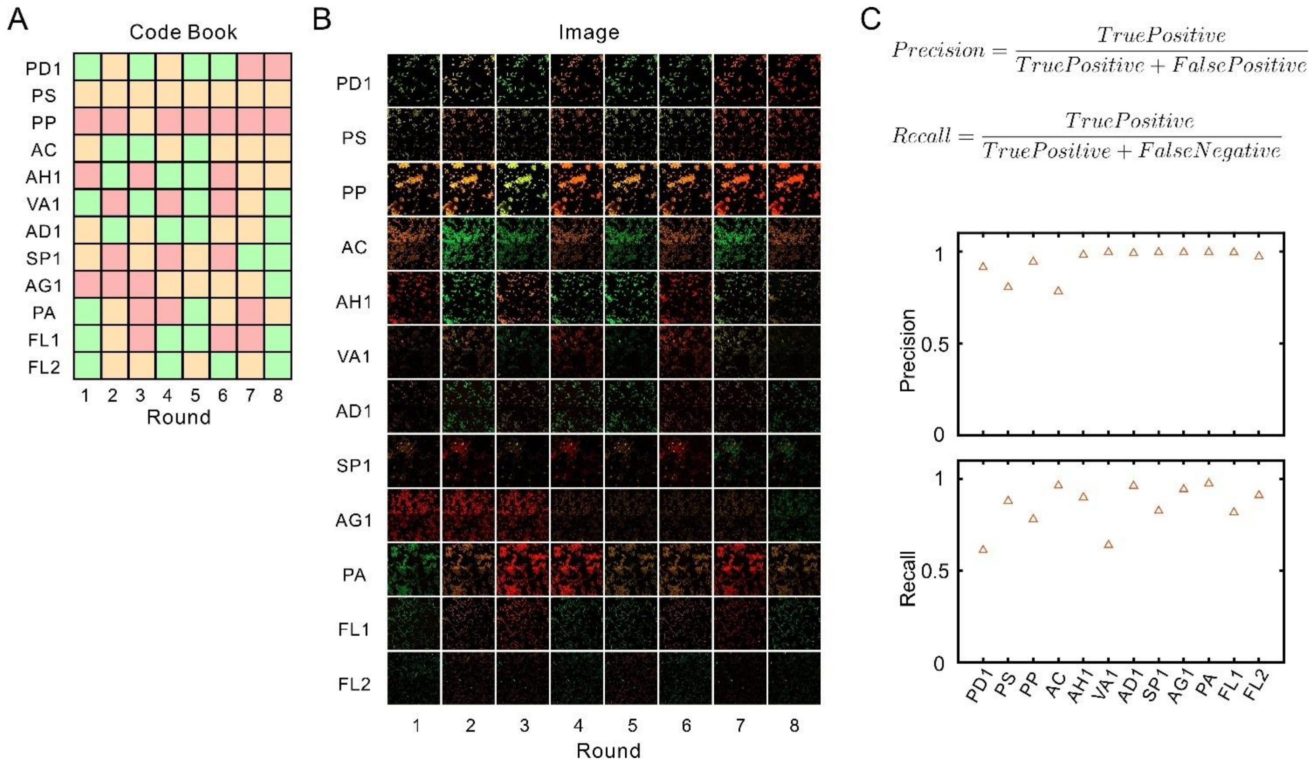


**Figure S5. Accuracy evaluation of SEER-FISH. A)** The codebook used for the experiment in Fig. 2C (R=8, HD≥4, S=12, F=3, Supplementary Table 5). The same codebook is also shown in Fig. 2A. **B)** The fluorescent images of each species during eight hybridization rounds. Fixed bacterial suspensions of 12 species were separately spotted onto 40-mm, #1.5 coverslip and air-dried. 8-round SEER-FISH was implemented according to the codebook in panel A. The fluorescence and phase contrast images of each species were acquired after each round of hybridization. **C)** Precision and recall of each bacterial species, based on identification of the 8-bit barcodes (Methods).

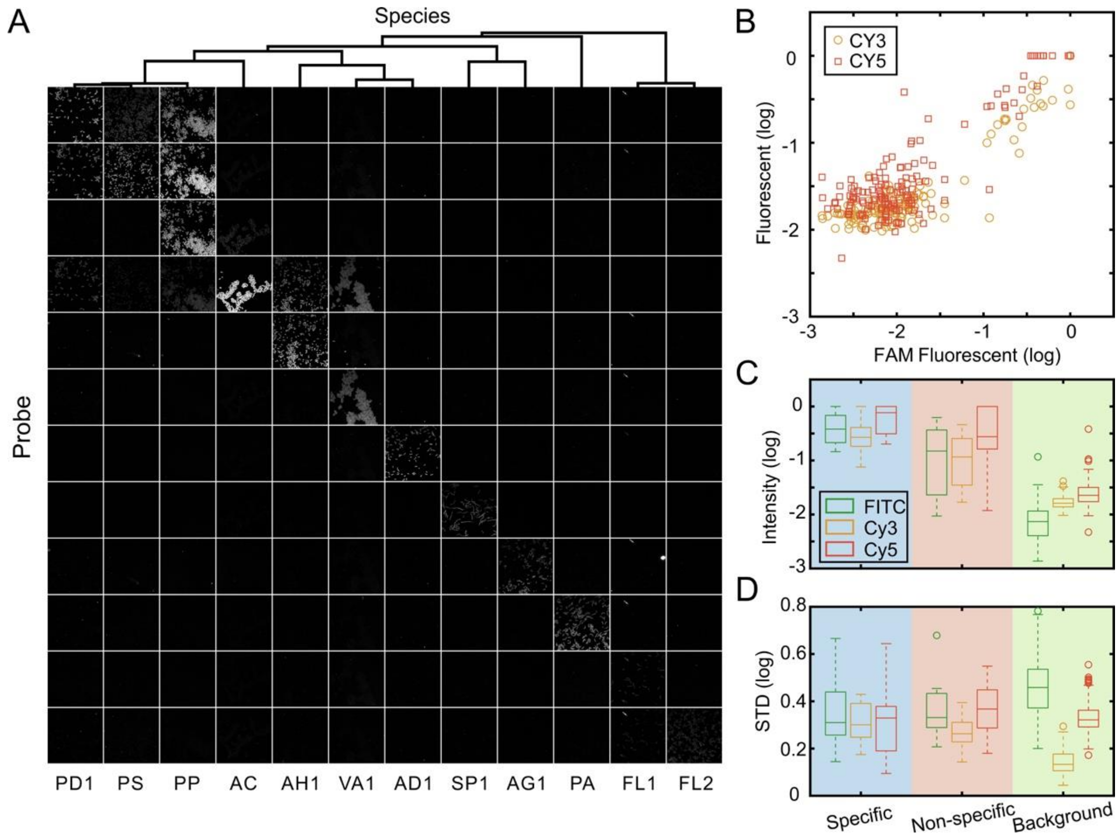


**Figure S6. Probe specificity analysis. A)** Fluorescent images of 12 pure bacterial cultures when hybridized with 12 candidate probes. 12 candidate probes were chosen to hybridize with pure cultures of target and non-target bacteria according to the codebook in Supplementary Table 5 (probe specificity analysis). To check the consistency of fluorescence signals between different fluorophores, 12 specific probes with three different conjugations (FAM, Cy3, Cy5) were all examined during 12 rounds of FISH imaging. The images of three fluorescence channels and phase-contrast images were collected for all species. **B)** The fluorescent intensity of probes conjugated to Cy3 or Cy5 is proportional to the intensity of probes conjugated to FAM. Pearson correlation is 0.88 (Cy3 to FAM, orange circles) and 0.90 (Cy5 to FAM, red squares), respectively. **C)** The mean fluorescent intensity in different channels. The probe-species pairs are grouped into 3 groups based on $\Delta G$ calculated by mathFISH. Three colored regions indicate specific binding ($\Delta G$ < -13.0 kcal/mol), non-specific binding (-7.9 kcal/mol > $\Delta G$ >-13.0 kcal/mol) and background ($\Delta G$ >-7.9 kcal/mol), respectively. The box plot indicates the interquartile range (25% to 75%) in each group. **D)** The standard deviation of fluorescent intensity in different channels.

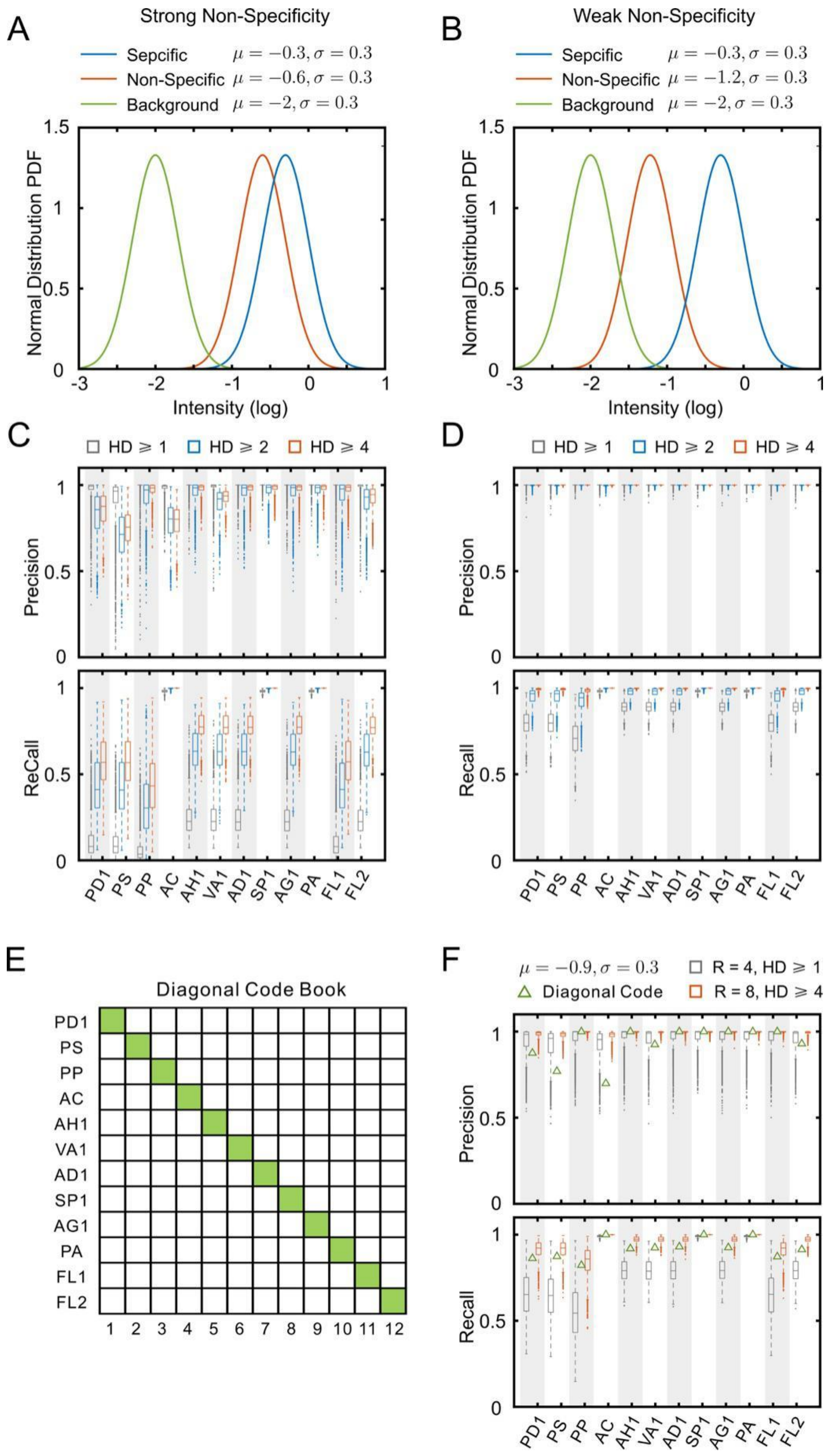


**Figure S7. Error-robust encoding enables improved accuracy. A) and B)** The mean intensity (log scale) of probe-species pairs from non-specific group is set as μ = -0.6 (A, strong non-specificity) and μ = -1.2 (B, weak non-specificity) respectively. **C) and D)** The precision and recall of 12 species obtained by simulations with different codebooks for the parameters used in A and B, respectively. The colored boxplot indicates the predicted distribution of precision and recall of SEER-FISH with 5000 randomly generated codebooks (F=3, R=8) with various minimal HD. Each box and the line inside the box labeled the first quartile, third quartile and median of each set of data. The length of the whiskers is set to 1.5 folds of the height of the box. The outliers are plotted as dots. Error-robust encoding schemes (HD≥2 and HD≥4) improve the overall performance of taxonomic identification. **E)** The diagonal codebook is a trivial codebook that uses N rounds of imaging to barcode N taxa. During each round, only one probe is present, and the barcode of each cell can be simply identified by finding the brightest round (without error-correction). **F)** The predicted precision and recall of taxon identification by exemplary coding schemes, including R4HD1, R8HD4 and the diagonal codebook. The boxplot indicates the predicted distribution of precision and recall of SEER-FISH with 5000 randomly generated codebooks (R4HD1, gray; R8HD4, red). The green triangles indicate the predicted precision and recall of the diagonal codebook illustrated in panel E. The mean intensity (log scale) of probe-species pairs for specific, non-specific and background hybridization is set as -0.3, -0.9 and -2, respectively.

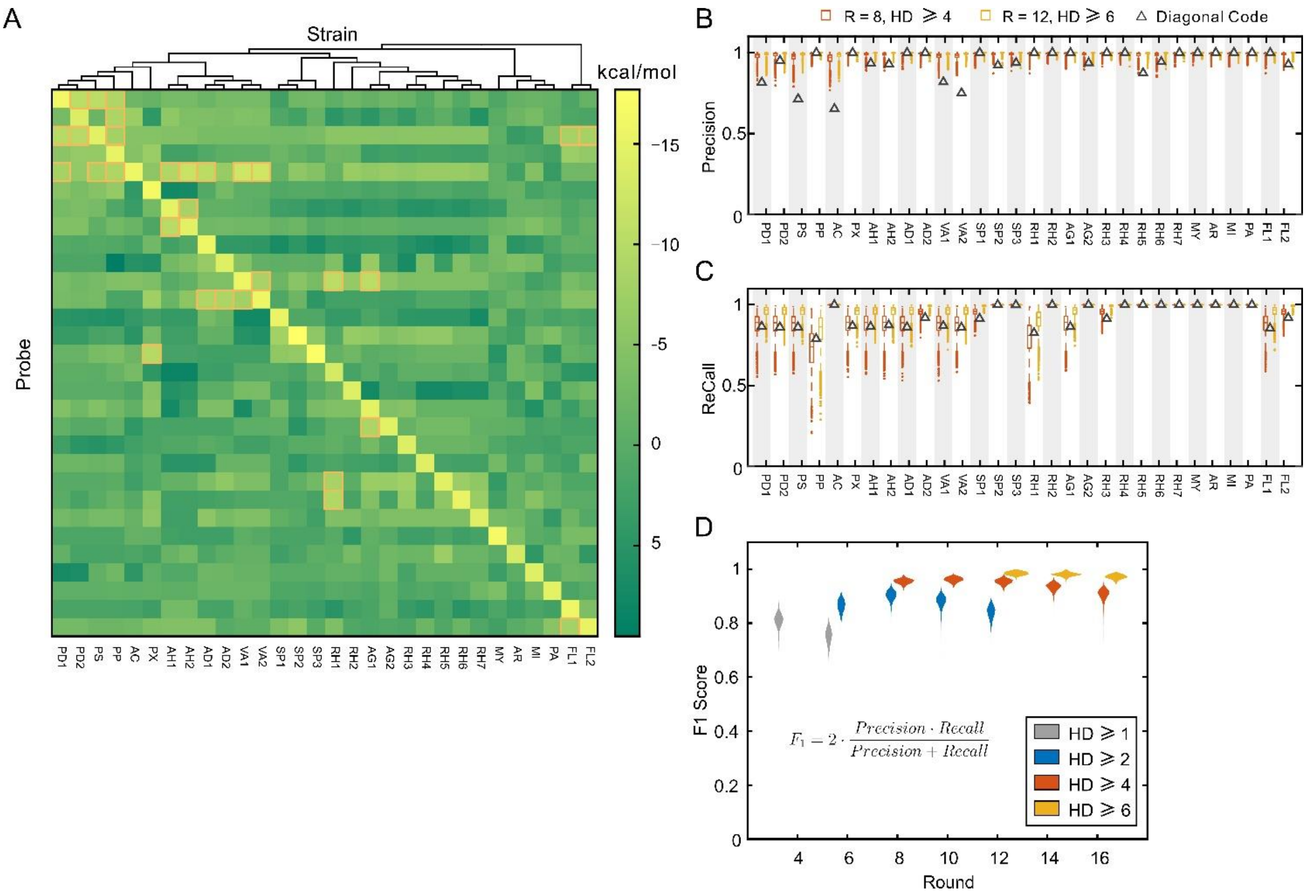


**Figure S8. Simulated performance of SEER-FISH for SynCom30. A)** The $\Delta G$ of each probe-specie pair was calculated by mathFISH. The pairs on the diagonal ($\Delta G$ <-13.0 kcal/mol) are grouped as specific binding. The pairs marked by orange boxes (-7.9 kcal/mol > $\Delta G$ >-13.0 kcal/mol) are grouped as non-specific binding. The rest ($\Delta G$>-7.9 kcal/mol) are grouped as background. **B) and C)** The precision and recall of 30 species obtained by simulation with 2 sets of 5000 randomly generated codebooks (Orange boxplot, F=3, R=8, HD≥4, S=30; Yellow boxplot, F=3, R=12, HD≥6, S=30) and the diagonal codebook (gray triangles). The diagonal code book is similar to the code book shown in Figure S7E (30 imaging rounds for 30 strains). The mean intensity (log scale) of probe-species pairs for specific, non-specific and background hybridization is set as -0.3, -0.9 and -2, respectively. **D)** Simulated performance of codebooks with varying levels of hybridization round (R, x-axis) and Hamming distance (HD, specified by colors). F1 score for 5000 randomly drawn codebooks (F=3, S=30) is shown. As expected, for a given HD, F1 score decreases with R; for a given R, F1 score increases with HD. The log-transformed fluorescent intensity ($\log(P_{pi})$) is drawn from a normal distribution with the mean fluorescence intensity of $10^{-0.3}$, $10^{-0.9}$ and $10^{-2}$ for specific binding ($\Delta G$ < -13.0 kcal/mol), non-specific binding (-7.9 kcal/mol > $\Delta G$ >-13.0 kcal/mol) and background ($\Delta G$ >-7.9 kcal/mol), respectively.

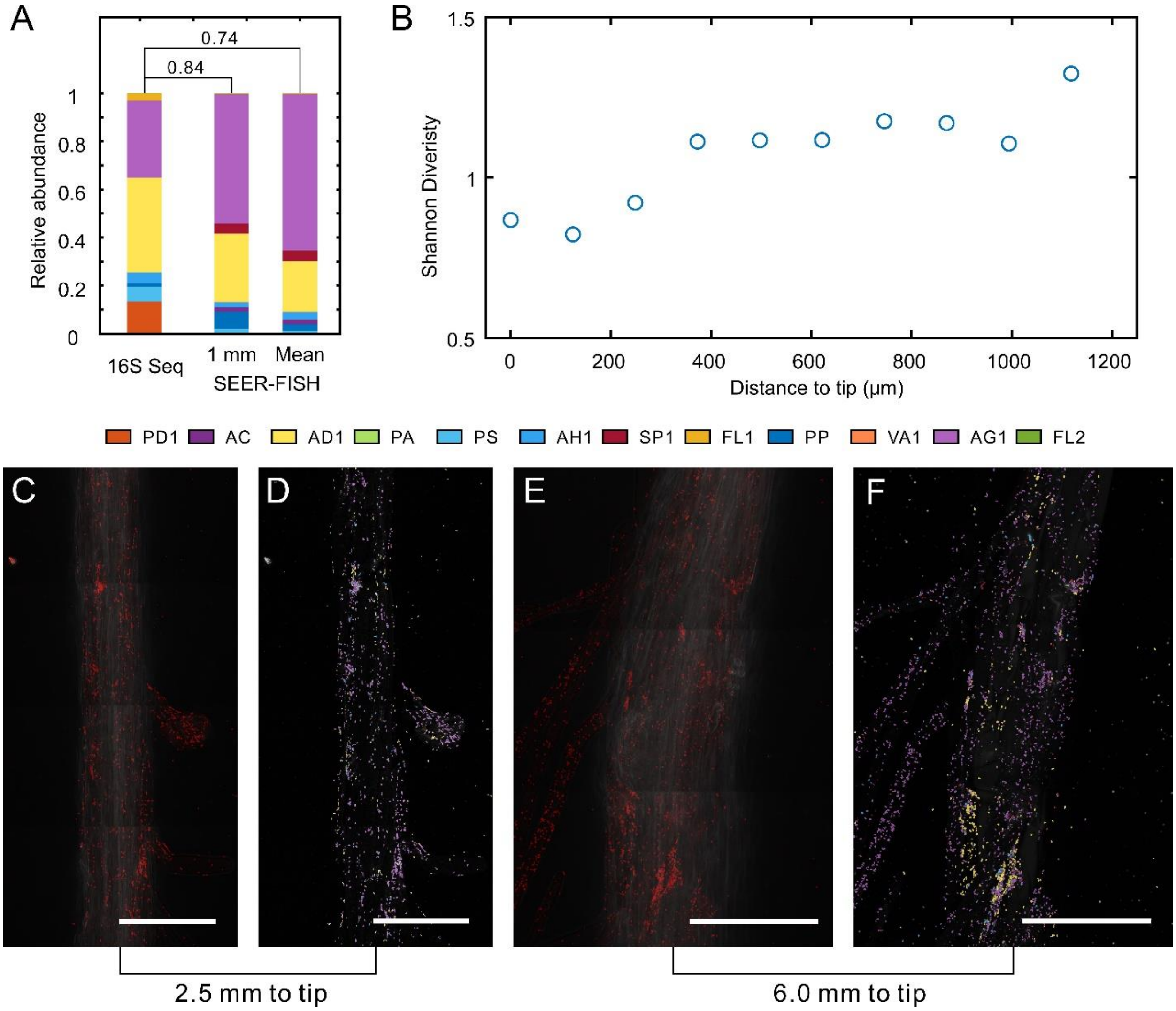


**Figure S9. Colonization of microbial communities on Arabidopsis root hairs. A)** The composition of the communities colonized on root measured by 16S amplicon sequencing and SEER-FISH. The Pearson correlation between the composition measured by sequencing and by SEER-FISH is indicated. 1 mm: the region within ~1mm from the root tip, as shown in Figure 4; Mean: averaged over 3 regions that are ~1 mm, 2.5 mm and 6 mm from the root tip. The difference in the composition profiles between 16S amplicon sequencing and imaging can be due to regional difference of measurements (average of ten whole roots by sequencing), various sources of bias in sequencing (e.g. DNA extraction, PCR bias), as well as in imaging (e.g. unidentified cells, sampling bias). **B)** The Shannon diversity of the communities within 1 mm of the root tip (Figure 4B). FOV = 125 μm（width）× 250 μm (length). **C-F)**. The 12-species synthetic community colonized on the root hair at 2.5 mm to the tip (C and D) and 6.0 mm to the tip (E and F). The bacteria cells are labeled by the universal probe in panel C and E. The bacteria cells are identified by SEER-FISH with 8 round barcodes (R=8, HD≥4, S=12, Supplementary Table 5) in panel D and F. Scale bar, 100 μm.

**Supplementary Table 1. Comparison of FISH-based methods for spatial mapping of microbial communities.**

| | SEER-FISH | HiPR-FISH | CLASI-FISH |
|---|---|---|---|
| **Multiplexity** | $F^N$ (F≥3, N≥20) | Up to 1023 | Up to 120 |
| **Error correction** | Yes | No | No |
| **Speed** | 20 min per round | 8-24 hours | 6-20 hours |
| **Probe costs** | Increase linearly with the number of target taxa and imaging rounds | Economic: 10 types of fluorescent readout probes | Two types of fluorescent probes for each target taxa |
| **Equipment needs** | Fluidics control for multi-round labeling | Spectral detector | Spectral detector |

**Supplementary Table 2. Information of bacterial strains used in this study.**

The whole genome sequences of root isolates were acquired from http://www.at-sphere.com/. 16S and 23S rRNA sequences of PP and PS were downloaded from NCBI. All 16S and 23S rRNA sequences are provided in Supplementary Files.

| Abb. | Strain ID | Phylum | Class | Family | Genus (species) |
|---|---|---|---|---|---|
| ***PD1** | root401 | *Proteobacteria* | *Gammaproteobacteria* | *Pseudomonadaceae* | *Pseudomonas* |
| **PD2** | root71 | *Proteobacteria* | *Gammaproteobacteria* | *Pseudomonadaceae* | *Pseudomonas* |
| ***PS** | WCS417 | *Proteobacteria* | *Gammaproteobacteria* | *Pseudomonadaceae* | *Pseudomonas (simiae)* |
| ***PP** | WCS358 | *Proteobacteria* | *Gammaproteobacteria* | *Pseudomonadaceae* | *Pseudomonas (putida)* |
| ***AC** | root1280 | *Proteobacteria* | *Gammaproteobacteria* | *Moraxellaceae* | *Acinetobacter* |
| **PX** | root630 | *Proteobacteria* | *Gammaproteobacteria* | *Xanthomonadaceae* | *Pseudoxanthomonas* |
| ***AH1** | root170 | *Proteobacteria* | *Betaproteobacteria* | *Alcaligenaceae* | *Achromobacter* |
| **AH2** | root83 | *Proteobacteria* | *Betaproteobacteria* | *Alcaligenaceae* | *Achromobacter* |
| ***AD1** | root70 | *Proteobacteria* | *Betaproteobacteria* | *Comamonadaceae* | *Acidovorax* |
| **AD2** | root217 | *Proteobacteria* | *Betaproteobacteria* | *Comamonadaceae* | *Acidovorax* |
| ***VA1** | root318D1 | *Proteobacteria* | *Betaproteobacteria* | *Comamonadaceae* | *Variovorax* |
| **VA2** | root473 | *Proteobacteria* | *Betaproteobacteria* | *Comamonadaceae* | *Variovorax* |
| ***SP1** | root241 | *Proteobacteria* | *Alphaproteobacteria* | *Sphingomonadaceae* | *Sphingomonas* |
| **SP2** | root1294 | *Proteobacteria* | *Alphaproteobacteria* | *Sphingomonadaceae* | *Sphingomonas* |
| **SP3** | root1497 | *Proteobacteria* | *Alphaproteobacteria* | *Sphingomonadaceae* | *Sphingopyxis* |
| **RH1** | root149 | *Proteobacteria* | *Alphaproteobacteria* | *Rhizobiaceae* | *Rhizobium* |
| **RH2** | root1203 | *Proteobacteria* | *Alphaproteobacteria* | *Rhizobiaceae* | *Rhizobium* |
| ***AG1** | root1240 | *Proteobacteria* | *Alphaproteobacteria* | *Rhizobiaceae* | *Agrobacterium* |
| **AG2** | Root491 | *Proteobacteria* | *Alphaproteobacteria* | *Rhizobiaceae* | *Agrobacterium* |
| **RH3** | root1298 | *Proteobacteria* | *Alphaproteobacteria* | *Rhizobiaceae* | *Rhizobium* |
| **RH4** | root483D2 | *Proteobacteria* | *Alphaproteobacteria* | *Rhizobiaceae* | *Rhizobium* |
| **RH5** | root1204 | *Proteobacteria* | *Alphaproteobacteria* | *Rhizobiaceae* | *Rhizobium* |
| **RH6** | root482 | *Proteobacteria* | *Alphaproteobacteria* | *Rhizobiaceae* | *Rhizobium* |
| **RH7** | root1212 | *Proteobacteria* | *Alphaproteobacteria* | *Rhizobiaceae* | *Rhizobium* |
| **MY** | root265 | *Actinobacteria* | *Actinobacteria* | *Mycobacteriaceae* | *Mycobacterium* |
| **AR** | root4 | *Actinobacteria* | *Actinobacteria* | *Microbacteriaceae* | *Agromyces* |
| **MI** | root166 | *Actinobacteria* | *Actinobacteria* | *Microbacteriaceae* | *Microbacterium* |
| ***PA** | root444D2 | *Firmicutes* | *Bacilli* | *Paenibacillaceae* | *Paenibacillus* |
| ***FL1** | Root186 | *Bacteroidetes* | *Flavobacteriia* | *Flavobacteriaceae* | *Flavobacterium* |
| ***FL2** | root901 | *Bacteroidetes* | *Flavobacteriia* | *Flavobacteriaceae* | *Flavobacterium* |

Abb.: abbreviation of strains

*Strains used for accuracy evaluation (Fig. 2C) and for SynCom12 and SynCom12_unequal (Figure 3-4).

All 30 strains were used for SynCom30 (Figure 3).

**Supplementary Table 3. Information of probes used in this study.**

All targeted probes designed in this study were conjugated to 3 types of fluorophores (FAM, Cy3 and Cy5). The universal probe (EUB338) was conjugated to Cy5.

| Abb. of target taxa | Target strain | Probe Sequence 5’-3’ | Target Region | $\Delta G$[a] （KJ/mol） | FA %[b] | 16S MM[c] | 23S MM[d] | WGS MM[e] |
|---|---|---|---|---|---|---|---|---|
| **PD1** | root401 | GTCCTACTCGATTTCACTTC | 23S | -16.3 | 31.1 | 7 | 3 | 3 |
| **PD2** | root71 | TCCGCCGCTGAATTCAGGAGC | 16S | -14 | 35.7 | 3 | 7 | 3 |
| **PS** | WCS417 | TTAGGTAACTGCCCTTCCTC | 16S | -14.7 | 34.7 | 2 | 7 | 2 |
| **PP** | WCS358 | CTGTGTCAGAGTTCCCGAAG | 16S | -15.3 | 30.8 | 3 | 7 | 3 |
| **AC** | root1280 | ACAAGTGATCCCCTGCTTTCC | 16S | -16 | 39.9 | 4 | 5 | 3 |
| **PX** | root630 | CATCTAATCGCGTGAGGCCTT | 16S | -16.6 | 37 | 6 | 7 | 3 |
| **AH1** | root170 | CCGGAACGTTTCTTTCCTGCC | 16S | -15.1 | 40.1 | 3 | 6 | 3 |
| **AH2** | root83 | CCATGACGTTTCTTTCCTGCC | 16S | -14.8 | 34.4 | 3 | 7 | 3 |
| **AD1** | root70 | CCCAGGTATTATCCAGAGTC | 16S | -16.1 | 34.7 | 5 | 6 | 3 |
| **AD2** | root217 | GTCATGGACCCCCTTTATT | 16S | -14.2 | 30.4 | 4 | 6 | 3 |
| **VA1** | root318D1 | TACCTTTCGGTGGGTTTCCC | 23S | -15.8 | 43 | 5 | 4 | 3 |
| **VA2** | root473 | GGTCGTTGTTAGCTGAAGCT | 23S | -16.1 | 35.6 | 6 | 3 | 3 |
| **SP1** | root241 | TGGTCTTTCGACATCATCCGG | 16S | -15.4 | 33.3 | 4 | 6 | 3 |
| **SP2** | root1294 | TCAACAGTCGTCCAGTGAGC | 16S | -17.7 | 38.2 | 4 | 6 | 3 |
| **SP3** | root1497 | TACTTGTCCAGTCAGTCGC | 16S | -17.4 | 39 | 4 | 5 | 3 |
| **RH1** | root149 | TCACACTCGCGTGCTCGCTG | 16S | -13.2 | 35.4 | 5 | 6 | 3 |
| **RH2** | root1203 | ATCTCTGCAAGTAGCCGGGC | 16S | -14.2 | 34.3 | 4 | 7 | 3 |
| **AG1** | root1240 | TCTCCGG TAACCGCGACCCA | 16S | -14.9 | 44.5 | 6 | 5 | 3 |
| **AG2** | Root491 | ACCCCGAATGTCAAGAGCTG | 16S | -14.7 | 27.9 | 4 | 6 | 3 |
| **RH3** | root1298 | AACGTCTCCGTAATCCGCGA | 16S | -14.1 | 30.9 | 3 | 7 | 3 |
| **RH4** | root483D2 | GCCGCTCGTATTGCTACGC | 16S | -14.3 | 37.3 | 5 | 4 | 2 |
| **RH5** | root1204 | CATTACTGCGTATCCTCAGCT | 23S | -14.6 | 29.9 | 7 | 3 | 3 |
| **RH6** | root482 | TTGCTCATGTATCCTCAGCT | 23S | -15.4 | 32.9 | 5 | 5 | 2 |
| **RH7** | root1212 | ACCTCTCGGTCGTATACGGTA | 16S | -14.4 | 31.2 | 4 | 6 | 3 |
| **MY** | root265 | GGCGCATGGTCATATTCGGT | 16S | -16.7 | 38.3 | 4 | 6 | 3 |
| **AR** | root4 | AGATGCCTCCGAGGGTCGT | 16S | -13.3 | 32.5 | 5 | 7 | 3 |
| **MI** | root166 | GCGGTCACGTCTCGTATCCA | 16S | -14.2 | 36.3 | 5 | 7 | 3 |
| **PA** | root444D2 | GGCCCATCTATAAGCCACAGA | 16S | -14.9 | 35.6 | 5 | 6 | 3 |
| **FL1** | Root186 | ACCGTCAAGTCCCGACACGT | 16S | -16 | 39.6 | 6 | 7 | 3 |
| **FL2** | root901 | GCGAGGTGGCTGCTCTCTGT | 16S | -15.2 | 43.6 | 3 | 7 | 3 |
| Universal | All | GCTGCCTCCCGTAGGAGT | 16S | N/A | N/A | 0 | N/A | N/A |

[a] $\Delta G$, the overall Gibbs free energy change for probe-target hybridization calculated by mathFISH.

[b] FA%, melting formamide concentration calculated by mathFISH.

[c] 16S MM, the minimal mismatch between the probe sequence and the 16S rRNA sequence of non-target strains.

Probes have at least three central mismatches to 16S rRNA sequences of non-target taxa, except the probe of PS (only two mismatches to PD1 and PD2).

[d] 23S MM, the minimal mismatch between the probe sequence and the 23S rRNA sequence of non-target strains.

[e] WGS MM, the minimal mismatch between the probe sequence and all mRNA sequences of non-target strains.

**Supplementary Table 4. Microscope setting used in this study.**

TD: transmitted detector. PMT: photomultiplier tube. HV: high voltage.

| Channel Setting | Excitation Wavelength | Laser Power | PMT HV | PMT Offset | Emission Wavelength | Filter Cube | Scan Mode | Scan Speed (µs/pixel) |
|---|---|---|---|---|---|---|---|---|
| FAM | 488 nm | 9 mW | 90 | 0 | 525 nm | 525/50 | Channel series | 0.5 |
| Cy3 | 561 nm | 3.2 mW | 55 | 0 | 595 nm | 595/50 | Channel series | 0.5 |
| Cy5 | 640 nm | 16 mW | 120 | 0 | 700 nm | 700/50 | Channel series | 0.5 |
| TD | N/A | N/A | 152 | 0 | N/A | N/A | Channel series | 0.5 |

**Supplementary Table 5. Codebooks used in this study.**

Color code equals 1, 2 and 3 for FAM, Cy3 and Cy5 fluorophores, respectively.

**R=8, HD≥4, S=12, F=3 (Figure 2, 3, 4 and S9)**

| Strain | Hybridization Rounds | | | | | | | |
|---|---|---|---|---|---|---|---|---|
| | **1** | **2** | **3** | **4** | **5** | **6** | **7** | **8** |
| **PD1** | 1 | 2 | 1 | 2 | 1 | 1 | 3 | 3 |
| **PS** | 2 | 2 | 2 | 2 | 2 | 2 | 2 | 2 |
| **PP** | 3 | 3 | 2 | 3 | 3 | 3 | 3 | 3 |
| **AC** | 2 | 1 | 1 | 2 | 1 | 2 | 2 | 2 |
| **AH1** | 3 | 1 | 3 | 1 | 1 | 3 | 2 | 2 |
| **VA1** | 1 | 3 | 1 | 3 | 1 | 3 | 2 | 1 |
| **AD1** | 2 | 1 | 2 | 1 | 1 | 2 | 2 | 1 |
| **SP1** | 2 | 3 | 2 | 3 | 2 | 3 | 1 | 1 |
| **AG1** | 3 | 3 | 3 | 2 | 2 | 2 | 2 | 1 |
| **PA** | 1 | 2 | 3 | 3 | 2 | 2 | 3 | 2 |
| **FL1** | 1 | 2 | 3 | 1 | 1 | 3 | 3 | 1 |
| **FL2** | 1 | 2 | 2 | 1 | 2 | 1 | 2 | 1 |

### R=8, HD≥4, S=30, F=3 (Figure 3)

| Strain | Hybridization Rounds | | | | | | | |
|---|---|---|---|---|---|---|---|---|
| | **1** | **2** | **3** | **4** | **5** | **6** | **7** | **8** |
| **PD1** | 1 | 3 | 2 | 1 | 3 | 2 | 2 | 3 |
| **PD2** | 1 | 1 | 1 | 1 | 3 | 3 | 3 | 3 |
| **PS** | 2 | 1 | 3 | 1 | 1 | 1 | 3 | 3 |
| **PP** | 1 | 1 | 3 | 2 | 3 | 1 | 3 | 1 |
| **AC** | 3 | 3 | 3 | 2 | 2 | 2 | 2 | 1 |
| **PX** | 2 | 2 | 2 | 3 | 2 | 1 | 3 | 3 |
| **AH1** | 1 | 3 | 1 | 3 | 3 | 1 | 3 | 2 |
| **AH2** | 1 | 2 | 3 | 3 | 2 | 2 | 3 | 2 |
| **AD1** | 2 | 2 | 2 | 2 | 2 | 2 | 2 | 2 |
| **AD2** | 2 | 1 | 1 | 2 | 1 | 2 | 1 | 2 |
| **VA1** | 3 | 1 | 2 | 3 | 3 | 1 | 2 | 1 |
| **VA2** | 3 | 2 | 1 | 3 | 1 | 2 | 2 | 3 |
| **SP1** | 1 | 2 | 1 | 2 | 2 | 3 | 1 | 2 |
| **SP2** | 2 | 2 | 2 | 2 | 1 | 1 | 1 | 1 |
| **SP3** | 3 | 2 | 3 | 3 | 2 | 1 | 1 | 1 |
| **RH1** | 3 | 1 | 2 | 1 | 3 | 2 | 3 | 2 |
| **RH2** | 1 | 2 | 3 | 1 | 3 | 1 | 1 | 3 |
| **AG1** | 3 | 1 | 3 | 1 | 1 | 3 | 2 | 2 |
| **AG2** | 2 | 3 | 1 | 1 | 1 | 3 | 1 | 3 |
| **RH3** | 1 | 1 | 2 | 2 | 1 | 1 | 2 | 2 |
| **RH4** | 1 | 1 | 2 | 3 | 3 | 3 | 1 | 2 |
| **RH5** | 1 | 2 | 2 | 1 | 1 | 2 | 1 | 2 |
| **RH6** | 2 | 3 | 1 | 2 | 3 | 3 | 3 | 1 |
| **RH7** | 3 | 1 | 2 | 2 | 1 | 3 | 3 | 1 |
| **MY** | 3 | 3 | 2 | 3 | 3 | 3 | 3 | 3 |
| **AR** | 2 | 2 | 1 | 1 | 1 | 1 | 2 | 2 |
| **MI** | 2 | 2 | 3 | 1 | 2 | 3 | 2 | 3 |
| **PA** | 1 | 3 | 1 | 3 | 1 | 3 | 2 | 1 |
| **FL1** | 2 | 1 | 1 | 3 | 3 | 1 | 1 | 3 |
| **FL2** | 2 | 1 | 1 | 3 | 2 | 3 | 3 | 2 |

### R=12, HD≥6, S=30, F=3 (Figure 3)

| Strain | Hybridization Rounds | | | | | | | | | | | |
|---|---|---|---|---|---|---|---|---|---|---|---|---|
| | 1 | 2 | 3 | 4 | 5 | 6 | 7 | 8 | 9 | 10 | 11 | 12 |
| PD1 | 1 | 2 | 1 | 1 | 1 | 3 | 3 | 1 | 1 | 2 | 3 | 2 |
| PD2 | 1 | 2 | 1 | 1 | 2 | 1 | 2 | 3 | 1 | 2 | 1 | 3 |
| PS | 2 | 1 | 1 | 1 | 2 | 3 | 3 | 2 | 2 | 3 | 2 | 3 |
| PP | 1 | 2 | 2 | 1 | 2 | 3 | 1 | 3 | 1 | 3 | 2 | 1 |
| AC | 3 | 3 | 3 | 3 | 3 | 2 | 1 | 1 | 1 | 3 | 2 | 3 |
| PX | 2 | 2 | 2 | 1 | 3 | 2 | 2 | 2 | 1 | 3 | 3 | 1 |
| AH1 | 2 | 3 | 2 | 2 | 1 | 2 | 3 | 3 | 3 | 1 | 1 | 1 |
| AH2 | 3 | 1 | 2 | 3 | 2 | 2 | 1 | 1 | 3 | 1 | 1 | 1 |
| AD1 | 2 | 1 | 2 | 2 | 3 | 2 | 1 | 2 | 3 | 2 | 2 | 1 |
| AD2 | 1 | 2 | 2 | 3 | 3 | 3 | 2 | 1 | 2 | 2 | 3 | 1 |
| VA1 | 2 | 1 | 3 | 1 | 1 | 1 | 2 | 2 | 1 | 3 | 1 | 3 |
| VA2 | 2 | 1 | 3 | 2 | 3 | 3 | 2 | 1 | 3 | 3 | 1 | 1 |
| SP1 | 1 | 1 | 1 | 2 | 2 | 1 | 3 | 1 | 3 | 2 | 2 | 1 |
| SP2 | 2 | 2 | 1 | 2 | 1 | 1 | 2 | 3 | 1 | 1 | 3 | 1 |
| SP3 | 1 | 2 | 1 | 1 | 1 | 2 | 3 | 2 | 2 | 1 | 1 | 3 |
| RH1 | 1 | 1 | 1 | 2 | 2 | 1 | 1 | 3 | 2 | 3 | 1 | 2 |
| RH2 | 2 | 1 | 1 | 1 | 2 | 3 | 2 | 3 | 3 | 2 | 3 | 2 |
| AG1 | 2 | 2 | 2 | 1 | 1 | 3 | 2 | 1 | 3 | 1 | 1 | 2 |
| AG2 | 2 | 2 | 1 | 3 | 1 | 3 | 3 | 3 | 3 | 1 | 2 | 3 |
| RH3 | 1 | 2 | 3 | 2 | 1 | 3 | 3 | 3 | 3 | 3 | 1 | 2 |
| RH4 | 3 | 2 | 2 | 1 | 1 | 1 | 1 | 2 | 3 | 3 | 3 | 2 |
| RH5 | 1 | 2 | 2 | 2 | 3 | 1 | 1 | 3 | 1 | 2 | 3 | 2 |
| RH6 | 3 | 1 | 3 | 2 | 2 | 2 | 3 | 1 | 2 | 3 | 1 | 3 |
| RH7 | 3 | 1 | 2 | 3 | 1 | 1 | 2 | 3 | 3 | 2 | 2 | 3 |
| MY | 1 | 2 | 2 | 2 | 2 | 2 | 3 | 2 | 2 | 2 | 2 | 2 |
| AR | 1 | 1 | 3 | 3 | 2 | 1 | 2 | 2 | 2 | 1 | 3 | 1 |
| MI | 2 | 2 | 1 | 2 | 3 | 2 | 1 | 2 | 1 | 1 | 1 | 2 |
| PA | 2 | 3 | 3 | 1 | 1 | 3 | 3 | 2 | 2 | 2 | 3 | 1 |
| FL1 | 2 | 1 | 3 | 1 | 2 | 1 | 1 | 3 | 2 | 2 | 2 | 1 |
| FL2 | 1 | 1 | 2 | 2 | 1 | 3 | 1 | 3 | 2 | 1 | 3 | 3 |

### Codebook used for probe specificity analysis (Figure S6)

| Strain | Hybridization Rounds | | | | | | | | | | | |
|---|---|---|---|---|---|---|---|---|---|---|---|---|
| | **1** | **2** | **3** | **4** | **5** | **6** | **7** | **8** | **9** | **10** | **11** | **12** |
| **PD1** | 1 | 0 | 0 | 0 | 2 | 0 | 0 | 0 | 3 | 0 | 0 | 0 |
| **PS** | 2 | 0 | 0 | 0 | 3 | 0 | 0 | 0 | 1 | 0 | 0 | 0 |
| **PP** | 3 | 0 | 0 | 0 | 1 | 0 | 0 | 0 | 2 | 0 | 0 | 0 |
| **AC** | 0 | 1 | 0 | 0 | 0 | 2 | 0 | 0 | 0 | 3 | 0 | 0 |
| **AH1** | 0 | 2 | 0 | 0 | 0 | 3 | 0 | 0 | 0 | 1 | 0 | 0 |
| **VA1** | 0 | 3 | 0 | 0 | 0 | 1 | 0 | 0 | 0 | 2 | 0 | 0 |
| **AD1** | 0 | 0 | 1 | 0 | 0 | 0 | 2 | 0 | 0 | 0 | 3 | 0 |
| **SP1** | 0 | 0 | 2 | 0 | 0 | 0 | 3 | 0 | 0 | 0 | 1 | 0 |
| **AG1** | 0 | 0 | 3 | 0 | 0 | 0 | 1 | 0 | 0 | 0 | 2 | 0 |
| **PA** | 0 | 0 | 0 | 1 | 0 | 0 | 0 | 2 | 0 | 0 | 0 | 3 |
| **FL1** | 0 | 0 | 0 | 2 | 0 | 0 | 0 | 3 | 0 | 0 | 0 | 1 |
| **FL2** | 0 | 0 | 0 | 3 | 0 | 0 | 0 | 1 | 0 | 0 | 0 | 2 |
